# Whole genome sequencing and the application of a SNP panel reveal primary evolutionary lineages and genomic variation in the lion (*Panthera leo*)

**DOI:** 10.1101/814103

**Authors:** L.D. Bertola, M. Vermaat, F. Lesilau, M. Chege, P.N. Tumenta, E.A. Sogbohossou, O.D. Schaap, H. Bauer, B.D. Patterson, P.A. White, H.H. de Iongh, J.F.J. Laros, K. Vrieling

## Abstract

**Background:** Previous phylogeographic studies of the lion (*Panthera leo*) have improved our insight into the distribution of genetic variation, as well as a revised taxonomy which now recognizes a northern (*Panthera leo leo*) and a southern (*Panthera leo melanochaita*) subspecies. However, existing whole range phylogeographic studies on lions either consist of very limited numbers of samples, or are focused on mitochondrial DNA and/or a limited set of microsatellites. The geographic extent of genetic lineages and their phylogenetic relationships remain uncertain, clouded by massive sampling gaps, sex-biased dispersal and incomplete lineage sorting.

**Results:** In this study we present results of low depth whole genome sequencing and subsequent variant calling in ten lions sampled throughout the geographic range, resulting in the discovery of >150,000 Single Nucleotide Polymorphisms (SNPs). Phylogenetic analyses revealed the same basal split between northern and southern populations as well as four population clusters on a more local scale. Further, we designed a SNP panel, including 125 autosomal and 14 mitochondrial SNPs, which was tested on >200 lions from across their range. Results allow us to assign individuals to one of these four major clades (West & Central Africa, India, East Africa, or Southern Africa) and delineate these clades in more detail.

**Conclusions:** The results presented here, particularly the validated SNP panel, have important applications, not only for studying populations on a local geographic scale, but also for tracing samples of unknown origin for forensic purposes, and for guiding conservation management of *ex situ* populations. Thus, these genomic resources not only contribute to our understanding of the evolutionary history of the lion, but may also play a crucial role in conservation efforts aimed at protecting the species in its full diversity.

## Background

Recent developments in next generation sequencing (NGS) techniques allow for the application of massive parallel sequencing to non-model organisms (1,2), such as the lion (*Panthera leo*). As a result, both the evolutionary history of a species and population histories can be reconstructed based on vastly expanded datasets (3,4). In addition to improved insight into the geographic distribution of genetic variation, this type of data can inform conservation efforts on how best to maintain this diversity (5,6). This is particularly relevant considering the dire situation of the lion populations in West and Central Africa, where downward populations trends are the strongest, and local extinctions have been reported in recent decades (7–11). The IUCN Red List currently classifies the lion as ‘Vulnerable’ across its range, but states that it would actually meet the criteria for ‘Endangered’ in East and Central Africa, with lions in West Africa being

‘Critically Endangered’ (12,13). Having a better understanding of intra-specific diversity in the lion can steer conservation efforts towards halting the loss of diversity.

The lion (*P. leo*) has been the subject of several phylogeographic studies which have provided insights into the evolution and distribution of genetic variation in the African populations (formerly subspecies *P. leo leo*) and its connection to the Indian population (formerly subspecies *P. leo persica*), located in and around the Gir National Park, Gujarat state, India. Although this Africa-Asia split has long been used to inform management (e.g. there is a separate studbook for Asiatic lions in zoos), this taxonomic distinction has since been overhauled. Phylogeographic studies played an important part in this, by providing improved understanding of the evolutionary history and relationships between populations. These studies included data from mitochondrial DNA (mtDNA) (14–23), autosomal DNA (18–20,22–24) and subtype variation in lion Feline Immunodeficiency Virus (FIV_Ple_) (19). Studies using mitochondrial markers and/or complete mitogenomes, describe a basal dichotomy, consisting of a northern group that includes populations from West and Central Africa as well as the Indian population (formerly recognized as a distinct subspecies), and a southern group with populations from East and Southern Africa (14,16,18,21).

Two studies have included Single Nucleotide Polymorphisms, although both have included only two sampling localities in West and Central Africa and exact sampling localities were often unknown, hence calling for improved sampling in the region (22,24). The distinction of lions in West and Central Africa was further corroborated by autosomal microsatellite data (20). Reliable inference of the evolutionary relationships between African and the Indian populations based on these data proved to be a challenge, as the extremely low variation in the Indian population led to an unresolved position in the distance tree (20).. Nevertheless, these newly described evolutionary relationships are reflected in a revised taxonomy which now recognizes a northern subspecies (*P. leo leo*) and a southern subspecies (*P. leo melanochaita*) (25).

However, each of the approaches used in the studies mentioned above have limitations. Although mtDNA is regarded as a useful genetic marker for gaining insight into phylogeographic patterns, partly because of its shorter coalescence time compared to nuclear markers, it may lead to ‘over-splitting’, reflecting fully coalesced groups based on mtDNA data, but incomplete lineage sorting of nuclear DNA (nuDNA) alleles. Moreover, mtDNA cannot identify admixture and represents only a single locus, potentially misrepresenting phylogenetic relationships due to the stochastic nature of the coalescent (26,27). In addition, sex-biased dispersal and gene flow will alter patterns derived from mitochondrial versus nuclear data. Because female lions exhibit strong philopatry whereas male lions are capable dispersers (28,29), phylogeographic patterns based on mtDNA in lions may overestimate divergence between populations. Inference of phylogenetic relationships from microsatellite data, on the other hand, is problematic due to their high variability and their mutation pattern leading to homoplasy. Moreover microsatellite studies typically employ only a few dozen markers which effectively represents <1% of the genome. Finally, most studies mentioned above were limited by sparse coverage of populations, notably in West and Central Africa. In particular, FIV_Ple_ prevalence is geographically restricted. Additional data from genome-wide markers, and covering more lion populations, are necessary to overcome these shortcomings. This will improve our understanding of the spatial distribution of variation in the lion, as well as help guide future conservation efforts that seek to preserve the species’ genetic diversity. As patterns of instra-specific variation are often shared across co-distributed taxa (21,30), a deeper insight into these patterns of lion variation will also be relevant for other species.

Here, we describe the discovery and phylogeographic analysis of genome-wide SNPs based on whole genomes and complete mitogenomes of ten lions, providing an overview of the intraspecific genomic variation. We further developed a SNP panel consisting of a subset (N=125) of the discovered SNPs, which was then used for genotyping >200 samples from 14 countries, representing almost the entire current distribution of the lion. This resolves phylogeographic breaks at a finer spatial resolution, and may serve as a reference dataset for future studies. Finally, we discuss the applications and future directions of high-throughput genotyping for wildlife research and conservation with recommendations on how they may contribute to future studies on conservation genomics.

## Results

### Sequencing

The sequencing runs yielded a total of 6.5·10^8^ reads for all samples combined, corresponding to an average coverage of ^~^3.8x per lion (see Supplemental Table 1 for coverage of autosomes and mitogenomes per individual). Following quality control, a total of 5.9·10^8^ reads (94.4%) were retained for subsequent alignment (Supplemental Table 1). More information on quality control is included in Supplemental Information 1 (and associated Supplemental Figures 1-4). Filtering of variable positions in lions with GATK yielded a total of 155,678 SNPs. Upon filtering for positions with coverage >3, we retained 118,270 SNPs of which 98,952 SNPs were called in at least five individuals (see Supplemental Table 2 for results for different levels of missing data). Missing data ranged from 90% (Benin) to 10% (Kenya) for all variable positions. Results reported for downstream phylogenetic analysis are based on the lion-specific variable positions which could be called in at least half of the included samples (98,952 SNPs).

### Whole genome data and complete mitogenomes

Phylogenetic analyses, based on 98,952 SNPs, show a well-supported dichotomy between the northern populations (Benin, Cameroon, Demoratic Republic of the Congo (DRC), India) and the southern ones (Somalia, Kenya, Zambia1, Zambia2, Republic of South Africa (RSA), Namibia) (Figure 1: left tree). Phylogenetic trees based on the mitochondrial genomes show the same basal dichotomy, with the Indian population nested within the northern populations (Figure 1: right tree), as was previously reported (21). However, it must be noted that the individual from RSA contains a haplotype from Namibia, likely the result of historic translocation, as was previously described (21). MrBayes, Garli and SVDquartets resulted largely in the same topology, although the three methods do not agree on the relationships of Zambia1, Zambia2, RSA and Namibia based on the autosomal SNPs. A dendrogram based on genotype likelihoods shows the same basal dichotomy, including a split within the Southern subspecies (Supplemental Information 2, Supplemental Figure 5).

**Figure 1.**
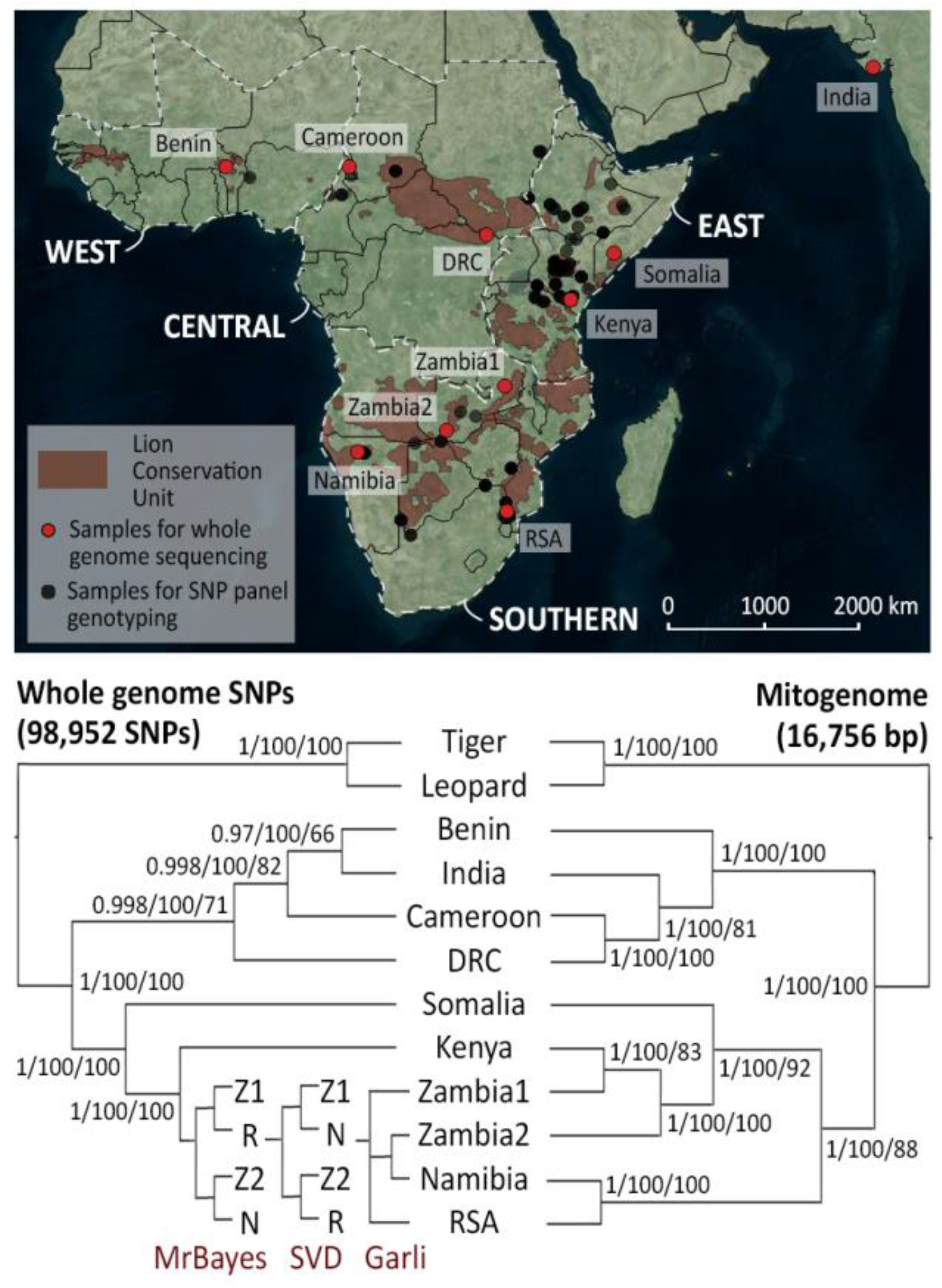
Distribution of lion sampling localities and inferred phylogenetic relationships between populations. Map indicating sampling locations of lions for whole genome sequencing (red) and SNP panel genotyping (black) (top). Red shading indicates Lion Conservation Units, and white delineation defines the regions (West, Central, East, and Southern), sensu the lion conservation strategies (93). Phylogenetic trees, based on 98,952 nuclear SNPs (left) and 16,756 bp mitogenomes (right) for 10 lions which were subjected to whole genome sequencing (bottom). Support values indicate posterior probabilities (MrBayes), bootstrap support from SVDquartets and bootstrap support from Garli. Topologies are indicated per method in the southern branch of the autosomal SNP tree, and each split is maximally supported, with the exception of the Zambia2+Namibia branch in the Garli tree, which received a bootstrap support of 96. Z1: Zambia1, Z2: Zambia2, N: Namibia, R: RSA.

PCA based on genotype likelihoods and called genotypes show a comparable pattern, with a strong geographic signal (Supplemental Information 2, Supplemental Figure 6). Clustering of the ten individuals using NGSadmix and sNMF, identifies the same basal split into a northern and a southern cluster (Figure 2A, sNMF results included in Supplemental Information 2, Supplemental Figure 7)). Based on the NGSadmix results, both Somalia and Kenya show signatures of admixture between the two subspecies. This is in line with results from the abbababa function in ANGSD. In the sNMF results, we observe a similar pattern for Somalia, but not for Kenya. This is further corroborated by the D-statistic (using GATK genotype calls) which resulted in significant admixture only for Somalia (but not for Kenya or DRC).

**Figure 2.**
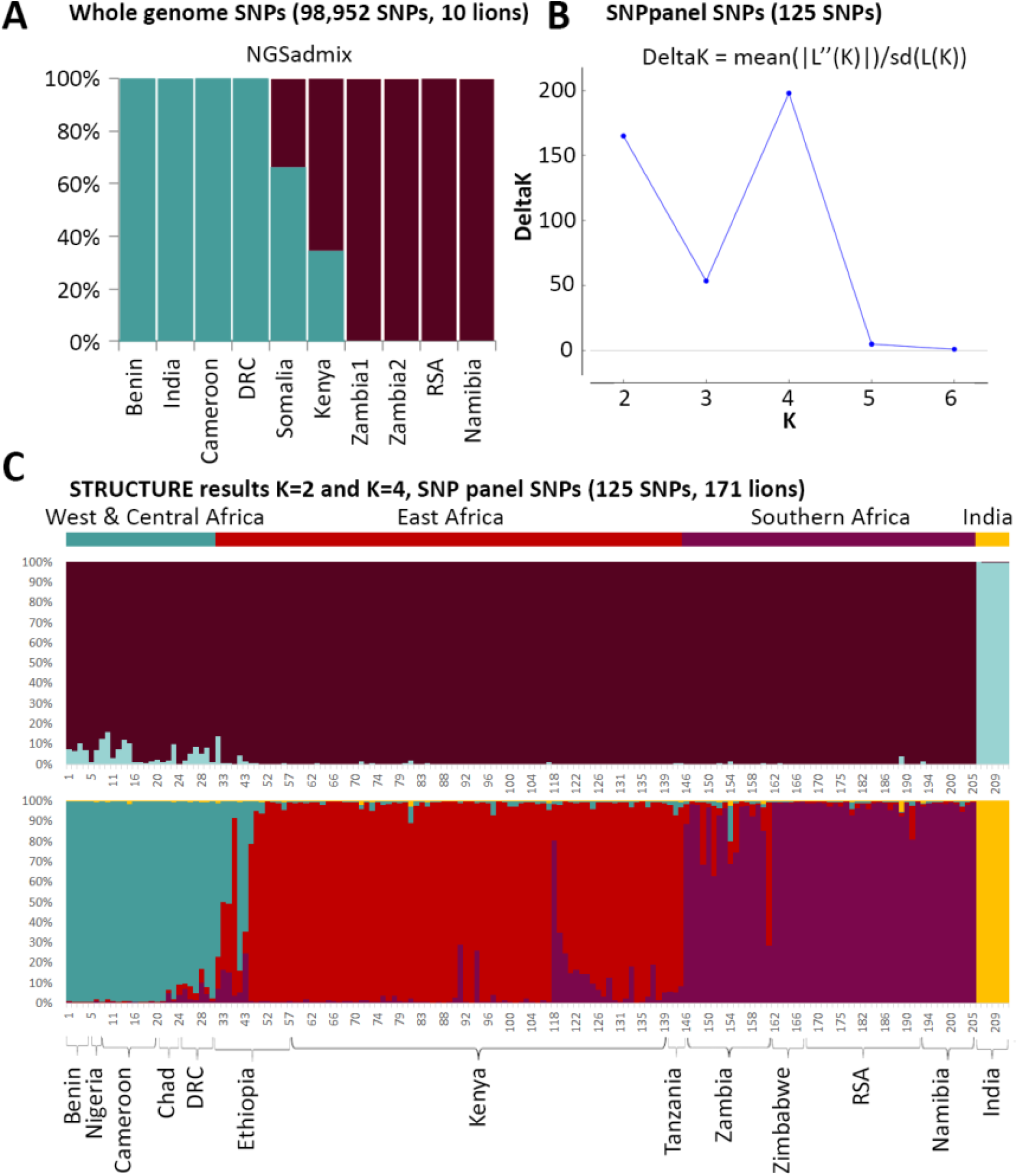
Assessment of population structure using NGSadmix and STRUCTURE. Results of NGSadmix (K=2) based on SNP genotype likelihoods from 10 lions (A). Result of STRUCTURE Harvester for a STRUCTURE run of 125 SNPs genotyped (B). Assignment values based on the STRUCTURE run for K=2 (upper bar plot) and K=4 (lower bar plot) for 171 lions (i.e. excluding samples with >25% missing data; see Supplemental Figure 6 for results for all 211 lions) (C). Sample numbers correspond with numbers in Supplemental Table 4.

### SNP panel data

A total of 211 samples from across the entire range of the lion were genotyped for 125 autosomal SNPs and 14 mitochondrial SNPs. Missing data for the autosomal SNPs ranged from to 0 to 115 with a median value of 11; for the mitochondrial data, these ranged from 0 to 10 with a median value of 0 (Supplemental Table 4, Supplemental Figure 8, included in Supplemental Information 2). As a confirmation of our initial SNP calling, the individuals which were subjected to whole genome sequencing have also been included in genotyping through the SNP panel. No contradictory genotypes were found, corroborating the results from our initial SNP calling. STRUCTURE suggests an optimal number of four clusters (Figure 2B) corresponding to West & Central Africa, India, East Africa, and Southern Africa (Figure 2C: lower bar plot). A lower peak can be detected for K=2, driven by India’s very low genetic diversity resulting in the formation of a strong separate cluster (Figure 2C: upper bar plot). Because missing data can affect the results of a STRUCTURE run, we present here the results excluding individuals with >25% missing data (N=171). STRUCTURE plots including the entire lion dataset (N=211) are available in Supplemental Figure 9 (included in Supplemental Information 2). Assignment values to clusters and to mitochondrial haplogroups are reported in Supplemental Table 5. In the PCA, African and Indian lions are distinguished as two clouds (Figure 3B). Removing the Indian population reveals more structure within African lions (Figure 3C), resulting in two nearly distinct clouds representing the northern (West & Central Africa) and the southern (East and Southern Africa) subspecies. PCA results based on whole genome genotype likelihoods are included as Figure 3A, showing a congruent geographic differentiation compared to the SNP panel SNPs. PCA results for the entire datasets (i.e. including samples with >25% missing data) are presented in Supplemental Information 2, Supplemental Figure 10.

**Figure 3.**
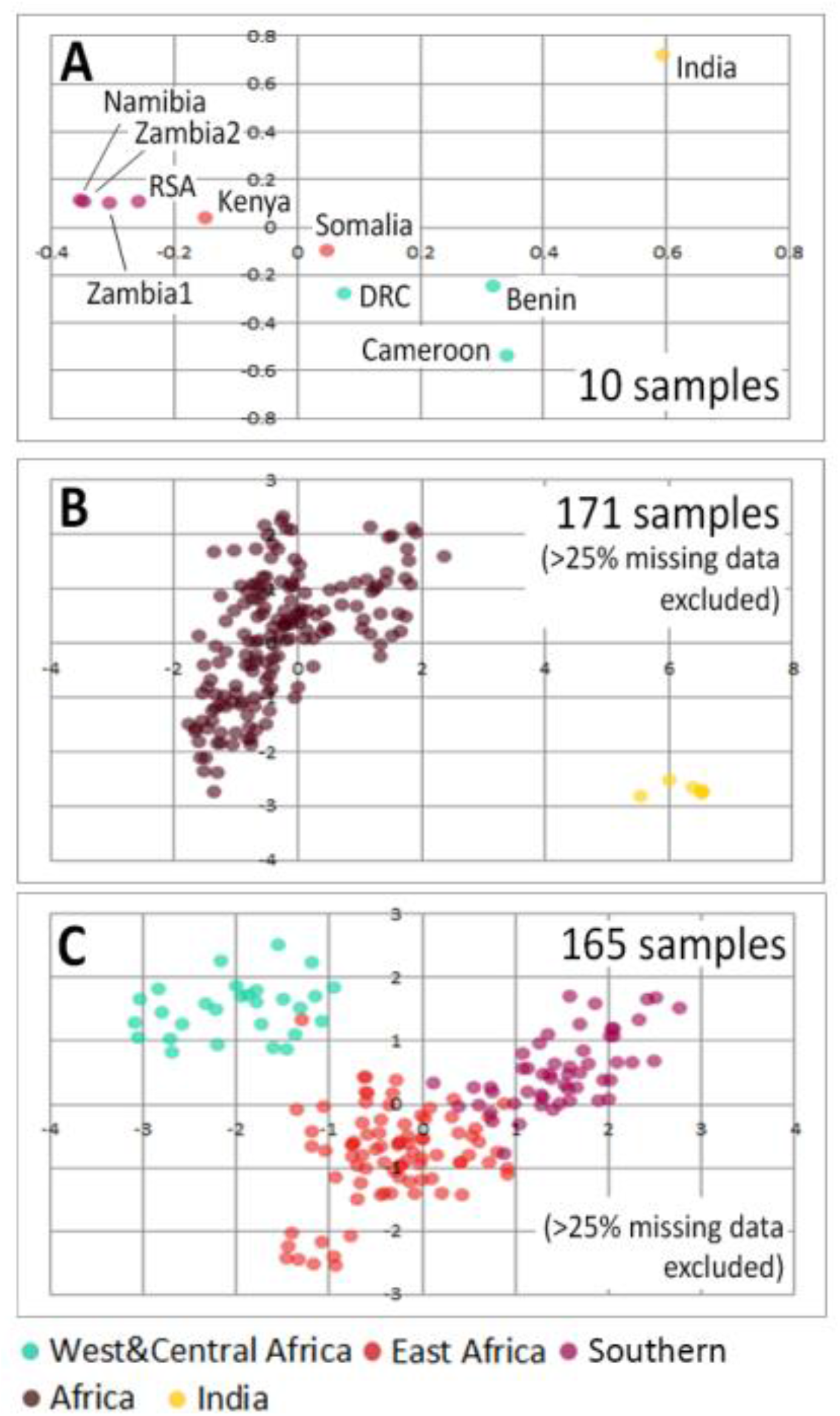
PCA plots based on whole genome genotype likelihoods and SNP panel SNPs. Plots are based on genotype likelihoods of 10 lions (A), 125 SNP panel SNPs in 171 lion samples (B), and after exclusion of India (C). Colours indicate region of origin.

EEMS infers the existence of corridors and barriers for dispersal and gene flow from the spatial decay of genetic similarity. Results show that the Central African rainforest is highlighted as a barrier (indicated with orange shading, Figure 4: upper row), whether or not the Indian population is included in the analysis. Barriers are further identified between East and Southern Africa, and across the Arabian peninsula. Diversity indices, which reflect the expected genetic dissimilarities of individuals sampled from the same deme, illustrate that the Indian population has a much lower genetic diversity than the African populations (Figure 4: bottom left). After repeating the analysis with only African populations, low genetic diversity is detected in West Africa and Southern Africa (Figure 4: bottom right).

**Figure 4.**
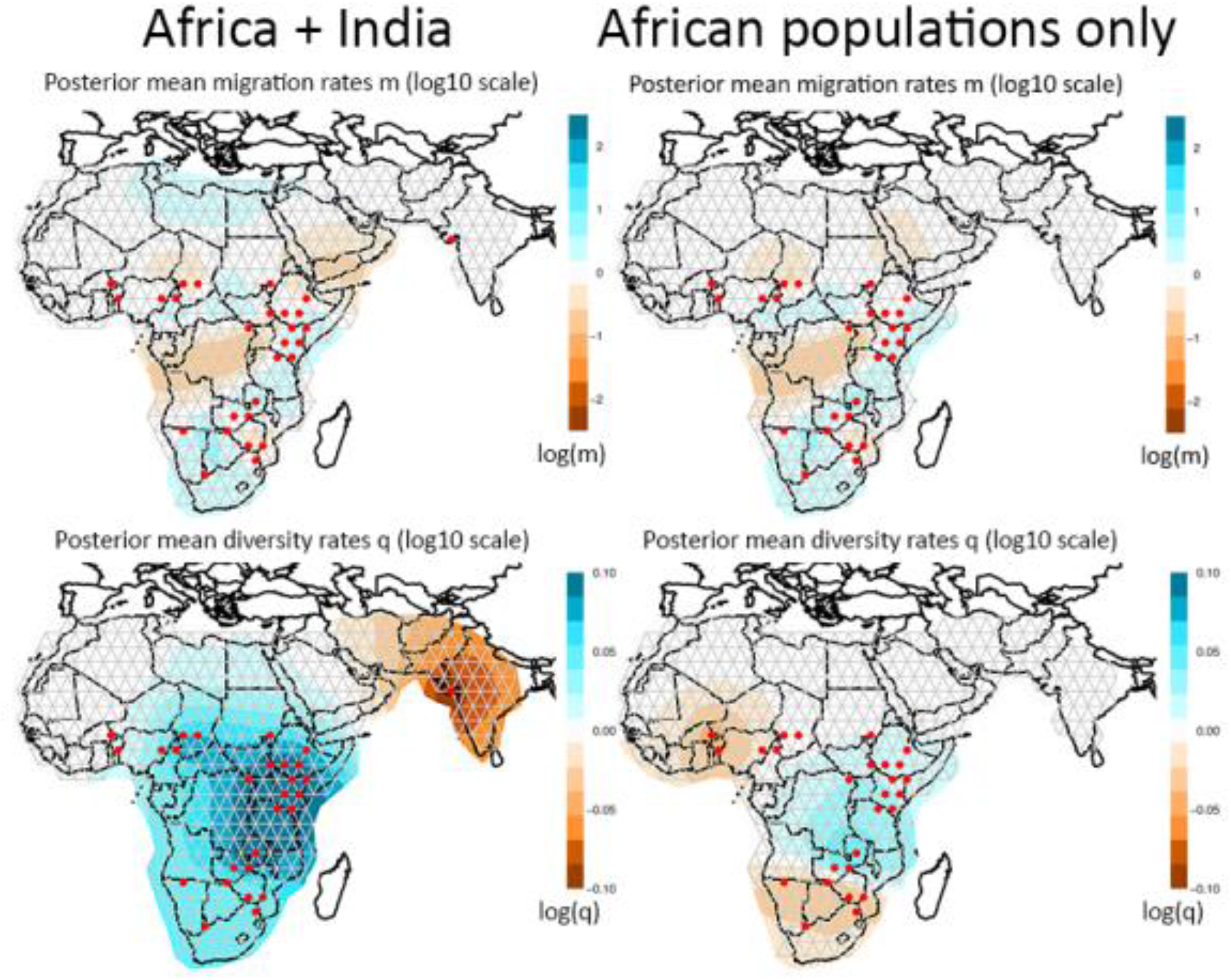
EEMS results based on 211 lions, including India (left), and 205 lions, excluding India (right). The upper row shows posterior mean migration rates, the bottom row shows posterior mean diversity rates. Orange colours indicate low values, blue colours indicate high values.

### Heterozygosity

Comparisons of levels of heterozygosity across datasets and to previously published data (18,20) (Supplemental Table 6) show that full genome observed heterozygosity is highly positively correlated with population-level observed heterozygosity derived from SNP panel SNPs and microsatellites (Supplemental Table 6: scatter plot and bar plot). The Indian population consistently shows a low observed heterozygosity for both SNPs and microsatellites (Supplemental Table 6: bar plot). However, for the estimates based on whole genome data, levels of heterozygosity are likely to be underestimated in the samples with low coverage (notably Benin).

## Discussion

### Whole genome data, complete mitogenomes and phylogeographic inference

Although SNPs can be powerful markers, ascertainment bias as a result of the study’s design is a concern (31). To reflect the full diversity of the species, it is necessary to base the SNP discovery, for SNPs which can subsequently used in a SNP panel, on samples representing all lineages. Different types of markers used in lion phylogeographic studies show largely congruent results with some local discrepancies (e.g. widespread East/Southern Africa haplogroup not recovered from nuDNA data (20) and admixture in the Kruger area (21,32)). Together, they provide useful criteria for selecting populations to be subjected to whole genome sequencing, as was done in this study

Based on the whole genome data, we explored both genotype likelihoods, derived from ANGSD and genotype calls derived from GATK, as well as different levels of missing data, balancing the number of SNPs and the number of samples with an accepted call at a given position. As a higher number of SNPs represents a denser sampling of the coalescent (e.g. see (33)), presented trees are based on SNPs which were present in at least 50% of the samples. Increasing the number of SNPs (and therefore also the amount of missing data), did not change the topology or the support of the phylogenetic trees. Simulation studies and studies using empirical data have shown that concatenated SNP data are able to produce reliable trees reflecting the true topology, as long as enough genes are sampled (34,35). The underlying assumption is that there is enough phylogenetic signal in the data, and that discordant coalescent histories will effectively cancel each other out when all histories are considered together. The conclusion that our concatenated phylogenetic trees (MrBayes and Garli) produce reliable topologies is supported by the fact that observed patterns are congruent with results of SVDquartets, which does assume multi-locus unlinked data. An exception is the more fine-scale topology for Southern Africa where patterns of differentiation are likely to be affected by continuous gene flow and incomplete lineage sorting. Further, described topologies are in line with previously described mitochondrial trees (14,21) with similar discordances as found in microsatellite datasets (20). Notably, the wide-spread haplogroup labeled as East/Southern Africa (21), stretching from Kenya to Namibia, is not recovered from microsatellite (20) or SNP data (this study). The previously mentioned dichotomy between the northern subspecies, *P. leo leo*, and the southern subspecies *P. leo melanochaita*, is also supported by the assignment values of NGSadmix and sNMF, with possible admixture where both subspecies overlap.

### SNP panel data

Based on the ten whole genomes, we generated a SNP panel of 125 autosomal and 14 mitochondrial SNPs which allows cost-effective genotyping of larger numbers of samples.

STRUCTURE results from the SNP panel data from lions across their natural range identify four clusters: West & Central Africa, India (the two clades of the northern subspecies, *P. leo leo*), East Africa, Southern Africa (the two clades of the southern subspecies, *P. leo melanochaita*) (Figure 2C). Main regions of admixture are Ethiopia and Zambia, as is visible in the STRUCTURE plots (Figure 2C), and was also found using microsatellite data (18,20) and mtDNA (36). It is well known that STRUCTURE is sensitive to groups of closely related individuals, such as siblings, family groups, or in our case, an inbred population (37,38) and that identification of ancestral populations may be an over-interpretation of the data, depending on demographic histories (39). Here, the West and Central African populations show an ancestral relationship to the Indian cluster, while also harbouring higher genetic diversity, leading to similarity to the southern subspecies cluster. The PCA results further illustrate that variation between African populations is masked by the extremely low diversity in the Indian population, likely the result of isolation and small population size due to multiple bottlenecks and subsequent genetic drift (also see results on heterozygosity, Supplemental Table 6). The variation between African populations only becomes apparent when exploring a higher number of clusuters (i.e., K=4; Figure 2C, bottom plot) or excluding the Indian populations (Figure 3C). After excluding the Indian population in the PCA, three overlapping clouds are apparent corresponding to West & Central Africa, East Africa and Southern Africa (Figure 3C). The individual which has a color code from East Africa (red) but falls in the West and Central Africa cluster (green) originates from Ethiopia, where both subspecies are known to overlap and admixture has been previously described (20,21). The distinction between East and Southern Africa seems to be more gradual in PCA space, which is in line with the widespread haplogroup that occurs throughout almost the entire region.

### Evolutionary history and connectivity

In order to put the phylogenetic patterns of the lion into an evolutionary perspective, it is worthwhile to explore current and historical barriers to lion dispersal. Paleoclimatic data show that cyclical contraction and expansion of vegetation zones, e.g. rain forest and desert, may have acted as temporal barriers to lion dispersal (21). The discrete genetic lineages recognizable in the mtDNA are likely to be the result of the restriction of suitable lion habitat to a number of refugia (21). The pattern found in mtDNA data of the lion is congruent with that of other African savanna species (21,30,40) and predicted refugial areas based on climate models (41). Faster coalescence times of mtDNA may have led to reciprocally monophyletic mtDNA clades in the lion, while isolation in refugia may not have lasted long enough for coalescence in autosomal markers (20,21). In addition, dispersal in lions is male-biased (28,29), which may explain the more discrete phylogeographic pattern found in mtDNA data. This is reflected by the fact that we do not retrieve a North East Africa cluster or a South West Africa cluster based on autosomal SNPs, even though they are represented by diverged mitochondrial haplotypes. Interestingly, the discrepancy in population structure between autosomal and mitochondrial markers in East and Southern Africa, where a wide-spread mitochondrial haplogroup occurs from Kenya to Namibia and autosomal markers suggest a phylogeographic break around Zambia and Mozambique, is identical to the discrepancy found in giraffe (42). Current barriers for gene flow seem to be mainly represented by the recent population disjunction in North Africa/Middle East and the longstanding barrier representing the Central African rain forest, as is also inferred by EEMS (Figure 4). Although the Rift valley, stretching from Ethiopia in the north to Malawi and Mozambique in the south, has been mentioned as a potential barrier for gene flow in the lion (Dubach *et al*. 2005; Barnett *et al*. 2006a; Barnett *et al*. 2006b; Bertola *et al*. 2011) and gene flow may be reduced in that region, co-occurrence of strongly diverged haplotypes and admixture detected in microsatellite and SNP data in Ethiopia indicates that the Rift valley does not represent an impenetrable barrier for lion dispersal (20,21).

### Applications for wildlife research and conservation

The design of a SNP panel and a reference dataset of >200 lions from 14 lion range states has important applications for wildlife research and conservation. First, it allows us to distinguish phylogeographic breaks and overlap between lineages which can be used to study the evolutionary history of the species in more detail, e.g. at the national level. This may have important conservation implications. For example, genomic data can be used to prioritize lion management at the population level, including providing key information for decisions regarding translocations of individuals within and among lion range states. Based on CITES documentation, >1000 live lions have been imported into lion range states since the 1980s with (future) potential to interbreed with resident lions (Bertola *et al*., in review). Human-mediated movement of wildlife, regarding lions and other species, only rarely take genetic considerations into account. Providing a baseline of the distribution of genetic variation and a tool to screen potential candidates for translocations, can help support these initiatives with scientific data. To assess the suitability of this SNP panel on a smaller geographic scale, we are currently genotyping >200 lion samples from across Kenya. The aim is to provide a robust tool for wildlife managers seeking genetic support for management decisions. Secondly, it enables tracing of lion samples of unknown origin, such as material confiscated from illegal trade chains or through anti-poaching efforts. Illegal trade in wildlife products is currently estimated to be the fifth largest illegal industry globally, and a major concern for conservationists (44–46). Part of the trade is thought to cater to domestic markets, but growing evidence suggests an increase in illegal shipments to international markets (47). Genetic toolkits can contribute to combatting wildlife crimes by identifying confiscated material, source populations and tracking trade routes (32,48–51). For lions, a tool that assists law enforcement intelligence has already been designed using mtDNA haplotypes (lionlocalizer.org), and it offers an ideal opportunity to include autosomal SNPs in future updates of this tool. Thirdly, a SNP panel can be used to guide breeding efforts for ex situ conservation (32,52). Recently, our SNP panel was used to genotype a batch of lion samples from institutions linked to the European Association for Zoos and Aquaria (EAZA). Based on the resulting information, managers are currently deciding how many and which lineages to include in future breeding efforts. In particular, given the dire situation for lions in West and Central Africa, and a severe under-representation of this lineage in captivity, the use of SNP data can be helpful to efficiently identify and conserve diversity *ex situ* and prevent inbreeding.

### Future perspectives

With the increase of genomic data alongside computational and technological developments, the field of conservation genomics is undergoing a major transition. Demographic histories and patterns of gene flow can be inferred in greater detail (53,54). Searching genomes for signatures of adaptation and deleterious alleles has been highlighted as a powerful tool for gauging the sensitivity of populations in a changing environment (55–59). New developments allowing SNP genotyping from poor quality (e.g., non-invasively collected) samples will further contribute to the applicability of SNP genotyping in the field (60). And, finally, the development of mobile, hand-held sequencing devices shows great promises for rapid, real-time identification of samples (61). Such tools represent tremendous potential for training and capacity building (62) especially for biodiverse countries with limited facilities to process samples, thereby making the field of wildlife research and conservation more democratized and inclusive.

### Conclusions

This study is the first to report phylogeographic relationships between lion populations throughout their entire range based on ^~^150,000 SNPs derived from whole genome sequencing. We present a SNP panel, containing 125 autosomal and 14 mitochondrial SNPs which has been validated on >200 individuals from 14 lion range states, spanning most of the lion’s range. The results reveal a detailed population structure and confirm a basal distinction between a northern and a southern subspecies that supports the recent revision of the lion’s taxonomy. We highlight several applications for the genomic resources we present here, and the SNP panel in particular. The samples on which the panel was tested can serve as a reference database for future research and conservation efforts.

## Methods

### Sampling

Blood or tissue samples of ten lions, representing the main phylogeographic groups as identified in previous studies (14,16,19–21) (Figure 1: map, Supplemental Table 1), were collected and preserved in a buffer solution (0.15 M NaCl, 0.05 M Tris-HCl, 0.001 M EDTA, pH = 7.5) at −20°C. All individuals included were either free-ranging lions or captive lions with proper documentation of their breeding history that included no known occurrences of hybridization between aforementioned lineages. A sample from an Amur leopard (*P. pardus orientalis*, captive) was included as an outgroup. The Amur tiger (*P. tigris altaica*) genome (63) was used as a reference for mapping of the lion and leopard reads. All samples were collected in full compliance with specific legally required permits (CITES and permits related to national legislation in the countries of origin). Details on laboratory protocols, sequencing, mapping to the reference genome, SNP calling, and quality control are given in Supplemental Information 1 (including Supplemental Figures 1-3).

### SNP calling and complete mitogenomes

After mapping of the reads to the reference genome (tiger reference genome from (63) and lion mitogenome from (21)), autosomal SNPs were called using two methods: 1) we used ANGSD (64) to derive genotype likelihoods, and 2) we called and filtered SNPs using GATK (65) and VCFtools (66) (details in Supplemental Information 1). Because these results stem from low coverage genomes, and acknowledging the potential confounding effect of missing data, we explored different levels of missing data applying the −minInd flag in ANGSD. For the SNPs derived from GATK we replaced positions with coverage <3 by an ambiguous nucleotide, and we applied 5 levels of filtering as follows: 1) all SNPs, 2) SNPs called in at least three samples, 3) SNPs called in at least five samples, 4) SNPs called in at least eight samples, and 5) SNPs called in all samples (Supplemental Table 2). Identified SNPs were attributed to a chromosome following the genomic architecture in the tiger (63) (Supplemental Table 2). Full mitogenomes were recovered by mapping all reads to a mitogenome reference generated with long range PCR and a known numt sequence, allowing us to differentiate between reads of mitochondrial and nuclear origin. This procedure is described in more detail in Supplemental Information 1 and (21).

Phylogenetic analyses were performed on the full mitogenomes and on the concatenated SNP datasets with varying levels of missing data with MrBayes v.3.1.2 (67,68) and Garli (69) using parameters as determined by MrModeltest2 (v.2.3) (70). MrBayes and Garli were run for one million generations and five million generations respectively, using a GTR substitution model with rate variation across sites set to equal. In addition, we ran SVDquartets (71), assuming multi-locus unlinked single-site data with 100 bootstrap replicates as implemented in PAUP* 4.0a164 (72). For the mitogenome data, the coalescent process in the model was disregarded. Nodes receiving >95% Posterior Probability (PP) in Bayesian analysis (MrBayes) and 0.7 bootstrap support in Maximum Likelihood (ML) analysis (Garli) and SVDquartets are considered to have significant support, as is common practice. A dendrogram was created on the genotype likelihoods derived from ANGSD, using the hclust algorithm. Genotype likelihoods (ANGSD) and genotype calls (GATK) were further subjected to Principal Component Analysis (PCA), using PCAngsd (73) and the PCA function from the Adegenet R package (74), respectively. Individual ancestry coefficients were estimated using NGSadmix (75) on the genotype likelihoods. For GATK genotypes, we used sparse non-negative matrix factorization algorithms in sNMF, as implemented in the R package LEA (76,77), exploring K=1-10, using 20 replicates and 4 values for the alpha regularization parameter (1, 10, 100 and 1,000). To explore putative admixture, we used the abbababa function in ANGSD and we calculated D-statistics (ABBA-BABA test) on the GATK genotypes using CalcD from the evobiR package (78) with 1000 bootstraps. We used Cameroon and Zambia1 as the ingroup as they are clearly assigned to *P. leo leo* and *P. leo melanochaita* respectively, and tiger as the outgroup.

### SNP panel data

In order to obtain better insight into the geographic locations of phylogeographic breaks, we made a selection of SNPs for inclusion in a SNP panel for genotyping more samples. We used the following criteria for the selection process: 1) minimum coverage of 20 for all lion samples combined, across 50 bp upstream and downstream of the SNP position, 2) maximum of one variable position in these 50 bp flanking regions, 3) high quality mapping of the flanking regions as identified by eye using IGV Genome Browser (79,80), 4) at least eight individuals represented at each selected site, 5) SNPs evenly spread across all chromosomes, as implied by the chromosomal architecture in tiger (max. one SNP per scaffold), 6) preferably both homozygotes and heterozygote genotype present among the ten genotyped lions (but sites for which each individual was scored as a heterozygote were excluded, as these may represent gene duplications). In addition, we included a total of 14 mitochondrial SNPs selected to represent each major branch within the mitochondrial phylogenetic tree (as described in (21)), as well as two diagnostic SNPs that represent the distinction between the northern and the southern subspecies. These mtDNA SNPs had already been assessed in a wide range of populations (21), making it more likely that the selected SNPs are diagnostic throughout the lion range. Finally, we ensured that the mitochondrial SNPs were not located in any of the nuclear copies (numts) which are known to exist in cats (81–85) by aligning our mitogenome sequences with known numt sequences. Coordinates of each of the SNPs and its associated chromosome, both for the tiger genome (63) and for a nearly-chromosome level lion genome (86) (lion genome was not available when the original mapping was done) are reported in Supplemental Table 3 and visualized in Supplemental Figure 4 (included in Supplemental Information 1). After test runs and quality control (see Supplemental Information 1), we retained 125 autosomal SNPs and 14 mtDNA SNPs which were used for genotyping 211 lions of known origin from 14 lion range states representing the entire geographic range of the species (Figure 1: map). There is a special focus on the region where the ranges of the northern and southern subspecies overlap i.e., Ethiopia and Kenya. This region is known to harbor haplotypes from strongly diverged lineages (21), and microsatellite data have suggested admixture (20). SNP genotyping was done at the SNP genotyping facility at Leiden University using allele-specific primers and KASP technology (LGC Genomics).

Resulting autosomal genotypes were analysed with STRUCTURE (87) using correlated allele frequencies and running the program for 5 million generations, discarding the first 500,000 generations as burn-in. Five replicates were run for K=1 to K=7. The optimal number of K was assessed by using the DeltaK method as implemented in STRUCTURE Harvester (88,89); CLUMPP (90) was used before generating the bar plots of population assignment. To reduce the effect of missing data on the assignment results, we ran STRUCTURE using the complete dataset (N=211) and a dataset excluding samples with >25% missing data (N=171). MtDNA SNPs were used to assign a specific haplotype to each individual, matching with previously described lineages (see delineation of haplogroups in (21)): West Africa, Central Africa, North East Africa, East/Southern Africa, South West Africa, and India.

A PCA was performed in Genalex (91), using the entire dataset (N=211) and the reduced dataset excluding samples with >25% missing data (N=177). Because the PCA is disproportionately affected by the very low genetic diversity of Indian lions, we repeatedthe analysis excluding this population, thereby allowing a more detailed picture from the African populations to emerge.

Patterns of connective zones and barriers were investigated using Estimated Effective Migration Surfaces (EEMS) (92) for all individuals genotyped with the SNP panel. Three independent runs were performed, using 10 million generations and discarding the first 5 million generations as burn-in. We followed the authors’ suggestions for tuning the proposal variances, using 0.1 and 1 for mSeedsProposalS2 and mEffctProposalS2 respectively, and 1.5 and 0.015 for qSeedsProposalS2 and qEffctProposalS2 respectively.

### Heterozygosity

The level of observed heterozygosity was assessed for each individual for which the whole genome had been sequenced, expressed as the proportion of heterozygote positions compared to the total number of SNPs, excluding positions with ambiguous nucleotides (i.e. positions which were not scored due to insufficient quality or coverage). This was done on the individual level for all SNPs, and on the populations level for all SNP panel SNPs which were successfully called in five or more individuals from the same population. Results were then compared to known levels of heterozygosity based on earlier studies using microsatellite data from the same populations (18,20).

## List of abbreviations

CITES: Convention on International Trade in Endangered Species of Wild Fauna and Flora
DRC: Demoratic Republic of the Congo
EAZA: European Association of Zoos and Aquaria
FIV_Ple_: Feline Immunodeficiency Virus
IUCN: International Union for Conservation of Nature
ML: Maximum Likelihood
mtDNA: mitochondrial DNA
NGS: Next Generation Sequencing
NP: National Park
nuDNA: nuclear DNA
PCA: Principal Component Analysis
RSA: Republic of South Africa
SNP: Single Nucleotide Polymorphism

## Declarations

### Ethics approval and consent to participate

Not applicable. All blood and tissue samples had been collected in the context of previous studies.

### Consent for publication

Not applicable

### Availability of data and materials

The datasets generated and analysed during the current study are available through NCBI under Bioproject ID … (accession numbers …-…) (fastq format), and included within the article and its supplementary information files (SNP panel genotypes).

### Authors’ contributions

L.D.B. performed analyses and wrote the manuscript, M.V. performed and J.F.J.L. supervised bioinformatics analyses, O.D.S. performed SNP genotyping, F.L., M.C., P.N.T., E.A.S., H.B., B.D.P. and P.A.W. supplied material and assisted with useful advice, H.H.d.I. and K.V. supervised analyses. All authors contributed to writing the manuscript.

## Acknowledgements

Samples were kindly provided by P. Henschel, Yohanna Saidu and the Nigerian National Park Service (Nigeria), S. Adam, R. Buij and B. Croes (Cameroon), N. Vanherle (CURESS, Chad), ICCN, Garamba NP (DRC), NABU (Ethiopia, Kafa BR), T. Jirmo (Kenya), South Africa National Parks (SANParks) (RSA), S. Miller, R. Groom, Save Valley Conservancy and DeBeers Venetia-Limpopo Nature Reserve/The Diamond Route (RSA and Zimbabwe), O. Aschenborn (Namibia), C.A.Driscoll (India), Safaripark Beekse Bergen (Hilvarenbeek, The Netherlands) (Somalia), Ouwehands Dierenpark (Rhenen, The Netherlands) (RSA), and Planckendael (Muizen, Belgium) (Leopard). We further thank N. Schidlo, R. Hennevelt, H. Buermans, Y. Ariyurek, and S. Greve-Onderwater for assisting in processing of the samples and BioIT for bioinformatics support.

## Supplemental Files

**Supplemental Information S1. Details on laboratory protocols, sequencing, mapping, SNP calling and quality control** (including Supplemental Figures S1-S4)

**Supplemental Information S2. Validation of phylogenetic inferences and populations structure from whole genome sequencing and SNP panel data** (including Supplemental Figures S5-S10)

**Supplemental Tables S1-S8 (see captions in Excel)**

## References

1. Ellegren H. Genome sequencing and population genomics in non-model organisms. Trends Ecol Evol [Internet]. 2014 Jan [cited 2014 Jul 10];29(1):51–63. Available from: http://www.ncbi.nlm.nih.gov/pubmed/24139972

2. Ekblom R, Galindo J. Applications of next generation sequencing in molecular ecology of non-model organisms. Heredity (Edinb) [Internet]. 2011 Jul [cited 2014 Jul 10];107(1):1–15. Available from: http://www.pubmedcentral.nih.gov/articlerender.fcgi?artid=3186121&tool=pmcentrez&rendertype=abstract

3. Shafer ABA, Wolf JBW, Alves PC, Bergström L, Bruford MW, Brännström I, et al. Genomics and the challenging translation into conservation practice. Trends Ecol Evol [Internet]. 2015 Dec [cited 2014 Dec 19];30(2):78–87. Available from: http://linkinghub.elsevier.com/retrieve/pii/S0169534714002511

4. Stephens ZD, Lee SY, Faghri F, Campbell RH, Zhai C, Efron MJ, et al. Big Data: Astronomical or Genomical? PLOS Biol [Internet]. 2015 Jul 7 [cited 2015 Jul 8];13(7):e1002195. Available from: http://dx.plos.org/10.1371/journal.pbio.1002195

5. Hendricks S, Anderson EC, Antao T, Bernatchez L, Forester BR, Garner B, et al. Recent advances in conservation and population genomics data analysis. Evol Appl. 2018;11(8):1197–211.

6. Supple MA, Shapiro B. Conservation of biodiversity in the genomics era. Genome Biol. 2018;19(1):1–12.

7. Craigie ID, Baillie JEM, Balmford A, Carbone C, Collen B, Green RE, et al. Large mammal population declines in Africa’s protected areas. Biol Conserv [Internet]. 2010 Sep [cited 2011 Jul 25];143(9):2221–8. Available from: http://linkinghub.elsevier.com/retrieve/pii/S0006320710002739

8. Bauer H, Chapron G, Nowell K, Henschel P, Funston P, Hunter LT, et al. Lion (Panthera leo) populations are declining rapidly across Africa, except in intesively managed areas. Proc Natl Acad Sci. 2015;1–6.

9. Riggio J, Jacobson A, Dollar L, Bauer H, Becker M, Dickman A, et al. The size of savannah Africa: a lion’s (Panthera leo) view. Biodivers Conserv [Internet]. 2012 Dec 2 [cited 2012 Dec 3];22(1):17–35. Available from: http://www.springerlink.com/index/10.1007/s10531-012-0381-4

10. Henschel P, Azani D, Burton C, Malanda GUY, Saidu Y, Sam M, et al. Lion status updates from five range countries in West and Central Africa. CATnews. 2010;52:34–9.

11. Brugière D, Chardonnet B, Scholte P. Large-scale extinction of large carnivores (lion Panthera leo, cheetah Acinonyx jubatus and wild dog Lycaon pictus) in protected areas of West and Central Africa. Trop Conserv Sci. 2015;8(2):513–27.

12. Bauer H, Packer C, Funston PF, Henschel P, Nowell K. The IUCN Red List of Threatened Species 2015: Panthera leo. 2015.

13. Henschel P, Coad L, Burton C, Chataigner B, Dunn A, MacDonald D, et al. The Lion in West Africa Is Critically Endangered. Hayward M, editor. PLoS One [Internet]. 2014 Jan 8 [cited 2014 Jan 9];9(1):e83500. Available from: http://dx.plos.org/10.1371/journal.pone.0083500

14. Bertola LD, van Hooft WF, Vrieling K, Uit de Weerd DR, York DS, Bauer H, et al. Genetic diversity, evolutionary history and implications for conservation of the lion (Panthera leo) in West and Central Africa. J Biogeogr [Internet]. 2011 Jul 31 [cited 2011 Sep 22];38(7):1356–67. Available from: http://doi.wiley.com/10.1111/j.1365-2699.2011.02500.x

15. Barnett R, Yamaguchi N, Barnes I, Cooper A. The origin, current diversity and future conservation of the modern lion (Panthera leo). Proc R Soc B Biol Sci [Internet]. 2006 Sep 7;273(1598):2119–25. Available from: http://www.ncbi.nlm.nih.gov/pubmed/16901830

16. Barnett R, Yamaguchi N, Shapiro B, Ho SY, Barnes I, Sabin R, et al. Revealing the maternal demographic history of Panthera leo using ancient DNA and a spatially explicit genealogical analysis. BMC Evol Biol [Internet]. 2014;14(1):70. Available from: http://www.biomedcentral.com/1471-2148/14/70

17. Dubach J, Patterson BD, Briggs MB, Venzke K, Flamand J, Stander P, et al. Molecular genetic variation across the southern and eastern geographic ranges of the African lion, Panthera leo. Conserv Genet [Internet]. 2005 Jan;6(1):15–24. Available from: http://www.springerlink.com/index/10.1007/s10592-004-7729-6

18. Dubach JM, Briggs MB, White PA, Ament BA, Patterson BD. Genetic perspectives on “Lion Conservation Units” in Eastern and Southern Africa. Conserv Genet [Internet]. 2013 Feb 3 [cited 2013 Feb 6];1942(Guggisberg 1961). Available from: http://link.springer.com/10.1007/s10592-013-0453-3

19. Antunes A, Troyer JL, Roelke ME, Pecon-Slattery J, Packer C, Winterbach C, et al. The evolutionary dynamics of the lion Panthera leo revealed by host and viral population genomics. PLoS Genet [Internet]. 2008 Nov;4(11):e1000251. Available from: http://www.ncbi.nlm.nih.gov/pubmed/18989457

20. Bertola LD, Tensen L, van Hooft P, White P a, Driscoll C a, Henschel P, et al. Autosomal and mtDNA Markers Affirm the Distinctiveness of Lions in West and Central Africa. PLoS One [Internet]. 2015 Jan [cited 2015 Oct 19];10(10):e0137975. Available from: http://www.ncbi.nlm.nih.gov/pubmed/26466139

21. Bertola LD, Jongbloed H, Van Der Gaag KJ, De Knijff P, Yamaguchi N, Hooghiemstra H, et al. Phylogeographic Patterns in Africa and High Resolution Delineation of Genetic Clades in the Lion (Panthera leo). Sci Rep [Internet]. 2016;6(August):1–11. Available from: http://dx.doi.org/10.1038/srep30807

22. Smitz N, Jouvenet O, Ligate FA, Crosmary W-G, Ikanda D, Chardonnet P, et al. A genome-wide data assessment of the African lion (Panthera leo) population genetic structure in Tanzania. PlosOne. 2018;1–24.

23. Curry CJ, Davis BW, Bertola LD, White PA, Murphy WJ. Spatiotemporal Genetic Diversity of Lions Reveals the Influence of Habitat Fragmentation Across Africa. Mol Biol Evol. 2020;1–27.

24. Manuel M De, Barnett R, Sandoval-Velasco M, Yamaguchi N, Vieira FG, Mendoza LZ, et al. The evolutionary history of extinct and living lions. Proc Natl Acad Sci. 2020;

25. Kitchener A, Breitenmoser-Würsten C, Eizirik E, Gentry A, Werdelin L, Wilting A, et al. A revised taxonomy of the Felidae. Cat News. 2017;80.

26. Zink RM, Barrowclough GF. Mitochondrial DNA under siege in avian phylogeography. Mol Ecol [Internet]. 2008 May [cited 2010 Jul 14];17(9):2107–21. Available from: http://www.ncbi.nlm.nih.gov/pubmed/18397219

27. Edwards S, Bensch S. Looking forwards or looking backwards in avian phylogeography? A comment on Zink and Barrowclough 2008. Mol Ecol [Internet]. 2009 Jul;18(14):2930–3; discussion 2934-6. Available from: http://www.ncbi.nlm.nih.gov/pubmed/19552688

28. Pusey AE, Packer C, Erhoff-Mulder MB. The Evolution of Sex-biased Dispersal in Lions. Behaviour. 1987;101(4):275–310.

29. Spong GF, Creel S. Deriving dispersal distances from genetic data. Proc Biol Sci [Internet]. 2001 Dec 22 [cited 2011 Feb 3];268(1485):2571–4. Available from: http://www.pubmedcentral.nih.gov/articlerender.fcgi?artid=1088917&tool=pmcentrez&rendertype=abstract

30. Lorenzen ED, Heller R, Siegismund HR. Comparative phylogeography of African savannah ungulates. Mol Ecol [Internet]. 2012 Aug [cited 2014 Feb 20];21(15):3656–70. Available from: http://www.ncbi.nlm.nih.gov/pubmed/22702960

31. Lachance J, Tishkoff SA. SNP ascertainment bias in population genetic analyses: Why it is important, and how to correct it. Bioessays. 2014;35(9):780–6.

32. Bertola LD. Genetic Diversity in the Lion (Panthera leo (Linnaeus 1758)): Unravelling the Past and Prospects for the Future. Leiden University; 2015.

33. Tripp EA, Tsai YHE, Zhuang Y, Dexter KG. RADseq dataset with 90% missing data fully resolves recent radiation of Petalidium (Acanthaceae) in the ultra-arid deserts of Namibia. Ecol Evol. 2017;7(19):7920–36.

34. Tonini J, Moore A, Stern D, Shcheglovitova M, Ort?? G. Concatenation and species tree methods exhibit statistically indistinguishable accuracy under a range of simulated conditions. PLoS Curr. 2015;7(TREE OF LIFE).

35. Xi Z, Liu L, Davis CC. The impact of missing data on species tree estimation. Mol Biol Evol. 2016;33(3):838–60.

36. Curry CJ, White PA, Derr JN. Mitochondrial Haplotype Diversity in Zambian Lions: Bridging a Gap in the Biogeography of an Iconic Species. PlosOne. 2015;1–14.

37. Waples RS, Anderson EC. Purging putative siblings from population genetic data sets: A cautionary view. Mol Ecol. 2017;1211–24.

38. Rodríguez-Ramilo ST, Wang J. The effect of close relatives on unsupervised Bayesian clustering algorithms in population genetic structure analysis. Mol Ecol Resour. 2012;12(5):873–84.

39. Lawson DJ, van Dorp L, Falush D. A tutorial on how not to over-interpret STRUCTURE and ADMIXTURE bar plots. Nat Commun [Internet]. 2018;9(1):1–11. Available from: http://dx.doi.org/10.1038/s41467-018-05257-7

40. Hewitt GM. The structure of biodiversity - insights from molecular phylogeography. Front Zool [Internet]. 2004 Oct 26 [cited 2014 Feb 26];1(1):4. Available from: http://www.pubmedcentral.nih.gov/articlerender.fcgi?artid=544936&tool=pmcentrez&rendertype=abstract

41. Levinsky I, Araújo MB, Nogués-Bravo D, Haywood AM, Valdes PJ, Rahbek C. Climate envelope models suggest spatio-temporal co-occurrence of refugia of African birds and mammals. Glob Ecol Biogeogr [Internet]. 2013 Mar 17 [cited 2014 Jun 7];22(3):351–63. Available from: http://doi.wiley.com/10.1111/geb.12045

42. Fennessy J, Bidon T, Reuss F, Vamberger M, Fritz U, Janke Correspondence A, et al. Multi-locus Analyses Reveal Four Giraffe Species Instead of One. Curr Biol [Internet]. 2016;26:1–7. Available from: http://dx.doi.org/10.1016/j.cub.2016.07.036

43. Barnett R, Yamaguchi N, Barnes I, Cooper A. Lost populations and preserving genetic diversity in the lion Panthera leo: Implications for its ex situ conservation. Conserv Genet [Internet]. 2006 Mar 1;7(4):507–14. Available from: http://www.springerlink.com/index/10.1007/s10592-005-9062-0

44. Nijman V, Morcatty T, Smith JH, Atoussi S, Shepherd CR, Siriwat P, et al. Illegal wildlife trade–surveying open animal markets and online platforms to understand the poaching of wild cats. Biodiversity [Internet]. 2019;20(1):58–61. Available from: https://doi.org/10.1080/14888386.2019.1568915

45. ‘t Sas-Rolfes M, Challender DWS, Hinsley A, Veríssimo D, Milner-Gulland EJ. Illegal Wildlife Trade: Patterns, Processes, and Governance. Annu Rev Environ Resour. 2019;44(1):1–28.

46. Biggs D, Cooney R, Roe D, Dublin HT, Allan JR, Challender DWS, et al. Developing a theory of change for a community-based response to illegal wildlife trade. Conserv Biol. 2017;31(1):5–12.

47. Williams VL, Loveridge AJ, Newton DJ, Macdonald DW. A roaring trade? The legal trade in Panthera leo bones from Africa to East-Southeast Asia. PLoS One. 2017;12(10):1–22.

48. Wasser S, Poole J, Lee P, Lindsay K, Dobson A, Hart J, et al. Elephants, Ivory, and Trade. Science (80-). 2010;327(March):1331–2.

49. Wasser SK, Brown L, Mailand C, Mondol S, Clark W, Laurie C, et al. Genetic assignment of large seizures of elephant ivory reveals major poaching hotspots. Science (80-). 2015;(June):1–7.

50. Wasser SK, Joseph Clark W, Drori O, Stephen Kisamo E, Mailand C, Mutayoba B, et al. Combating the illegal trade in African elephant ivory with DNA forensics. Conserv Biol [Internet]. 2008 Aug [cited 2010 Aug 30];22(4):1065–71. Available from: http://www.ncbi.nlm.nih.gov/pubmed/18786100

51. Mondol S, Sridhar V, Yadav P, Gubbi S, Ramakrishnan U. Tracing the Geographic Origin of Traded Leopard Body Parts in the Indian Subcontinent with DNA-Based Assignment Tests. Conserv Biol [Internet]. 2014 Nov 5 [cited 2014 Nov 11];00(0):1–9. Available from: http://www.ncbi.nlm.nih.gov/pubmed/25376464

52. Henry P, Miquelle D, Sugimoto T, McCullough DR, Caccone A, Russello M a. In situ population structure and ex situ representation of the endangered Amur tiger. Mol Ecol [Internet]. 2009 Aug [cited 2013 Oct 30];18(15):3173–84. Available from: http://www.ncbi.nlm.nih.gov/pubmed/19555412

53. Kamm JA, Terhorst J, Durbin R, Song YS. Efficiently inferring the demographic history of many populations with allele count data. bioRxiv [Internet]. 2018;287268. Available from: https://www.biorxiv.org/content/early/2018/03/23/287268

54. Elleouet JS, Aitken SN. Exploring Approximate Bayesian Computation for inferring recent demographic history with genomic markers in nonmodel species. Mol Ecol Resour. 2018;18(3):525–40.

55. Harrisson KA, Pavlova A, Telonis-Scott M, Sunnucks P. Using genomics to characterize evolutionary potential for conservation of wild populations. Evol Appl [Internet]. 2014 Mar 14 [cited 2014 Sep 5];7(9):1008–25. Available from: http://doi.wiley.com/10.1111/eva.12149

56. Funk WC, Mckay JK, Hohenlohe PA, Allendorf FW. Harnessing genomics for delineating conservation units. Trends Ecol Evol [Internet]. 2012 Sep [cited 2014 Jul 10];27(9):489–96. Available from: http://dx.doi.org/10.1016/j.tree.2012.05.012

57. Funk W, Lovich R, Hohenlohe P, Hofman C, Morrison S, Sillett T, et al. Adaptive divergence despite strong genetic drift: genomic analysis of the evolutionary mechanisms causing genetic differentiation in the island fox (Urocyon littoralis). Mol Ecol. 2016;25(10):2176–94.

58. Razgour O, Forester B, Taggart JB, Bekaert M, Juste J, Ibáñez C, et al. Considering adaptive genetic variation in climate change vulnerability assessment reduces species range loss projections. Proc Natl Acad Sci U S A. 2019;116(21):10418–23.

59. Henn BM, Botigué LR, Bustamante CD, Clark AG, Gravel S. Estimating the mutation load in human genomes. Nat Rev Genet. 2015;16(6):333–43.

60. Natesh M, Taylor RW, Truelove NK, Hadly EA, Palumbi SR, Petrov DA, et al. Empowering conservation practice with efficient and economical genotyping from poor quality samples. Methods Ecol Evol. 2019;2019(August 2018):1–7.

61. Johri S, Solanki J, Cantu VA, Fellows SR, Edwards RA, Moreno I, et al. ‘Genome skimming’ with the MinION hand-held sequencer identifies CITES-listed shark species in India’s exports market. Sci Rep [Internet]. 2019;9(1):1–13. Available from: http://dx.doi.org/10.1038/s41598-019-40940-9

62. Pomerantz A, Peñafiel N, Arteaga A, Bustamante L, Pichardo F, Coloma LA, et al. Real-time DNA barcoding in a rainforest using nanopore sequencing: Opportunities for rapid biodiversity assessments and local capacity building. Gigascience. 2018;7(4):1–14.

63. Cho YS, Hu L, Hou H, Lee H, Xu J, Kwon S, et al. The tiger genome and comparative analysis with lion and snow leopard genomes. Nat Commun. 2013;4(2433).

64. Korneliussen TS, Albrechtsen A, Nielsen R. ANGSD: Analysis of Next Generation Sequencing Data. BMC Bioinformatics. 2014;15(1):1–13.

65. McKenna A, Hanna M, Banks E, Sivachenko A, Cibulskis K, Kernytsky A, et al. The Genome Analysis Toolkit: A MapReduce framework for analyzing next-generation DNA sequencing data. Genome Res. 2010;20:1297–303.

66. Danecek P, Auton A, Abecasis G, Albers CA, Banks E, DePristo MA, et al. The variant call format and VCFtools. Bioinformatics. 2011 Aug;27(15):2156–8.

67. Huelsenbeck JP, Ronquist F. MRBAYES: Bayesian inference of phylogenetic trees. Bioinformatics [Internet]. 2001 Aug;17(8):754–5. Available from: http://www.ncbi.nlm.nih.gov/pubmed/11524383

68. Ronquist F, Teslenko M, van der Mark P, Ayres DL, Darling A, Höhna S, et al. MrBayes 3.2: efficient Bayesian phylogenetic inference and model choice across a large model space. Syst Biol [Internet]. 2012 May [cited 2013 Feb 28];61(3):539–42. Available from: http://www.pubmedcentral.nih.gov/articlerender.fcgi?artid=3329765&tool=pmcentrez&rendertype=abstract

69. Zwickl DJ. Genetic algorithm approaches for the phylogenetic analysis of large biological sequence datasets under the maximum likelihood criterion. The University of Texas at Austin; 2006.

70. Nylander JAA. MrModeltest v2.3. Uppsala, Sweden: Evolutionary Biology Centre, Uppsala University; 2004. p. Program distributed by the author.

71. Chifman J, Kubatko L. Quartet inference from SNP data under the coalescent model. Bioinformatics. 2014;30(23):3317–24.

72. Swofford DL. PAUP*: Phylogenetic analysis using parsimony (*and other methods), version 4b10. Sinauer, Sunderland, Massachusetts, USA.; 2002.

73. Meisner J, Albrechtsen A. Inferring population structure and admixture proportions in low-depth NGS data. Genetics. 2018;210(2):719–31.

74. Jombart T. Adegenet: A R package for the multivariate analysis of genetic markers. Bioinformatics. 2008;24(11):1403–5.

75. Skotte L, Korneliussen TS, Albrechtsen A. Estimating individual admixture proportions from next generation sequencing data. Genetics. 2013;195(3):693–702.

76. Frichot E, François O. LEA: An R package for landscape and ecological association studies. Methods Ecol Evol. 2015;6(8):925–9.

77. Frichot E, Mathieu F, Trouillon T, Bouchard G, François O. Fast and efficient estimation of individual ancestry coefficients. Genetics. 2014;196(4):973–83.

78. Durand EY, Patterson N, Reich D, Slatkin M. Testing for ancient admixture between closely related populations. Mol Biol Evol [Internet]. 2011 Aug [cited 2014 May 25];28(8):2239–52. Available from: http://www.pubmedcentral.nih.gov/articlerender.fcgi?artid=3144383&tool=pmcentrez&rendertype=abstract

79. Robinson JT, THorvaldsdóttir H, Winckler W, Guttman M, Lander ES, Getz G, et al. Integrative Genomic Viewer. Nat Biotechnol [Internet]. 2013;29(1):24–6. Available from: https://www.broadinstitute.org/igv/node/250

80. Thorvaldsdóttir H, Robinson JT, Mesirov JP. Integrative Genomics Viewer (IGV): High-performance genomics data visualization and exploration. Brief Bioinform. 2013;14(2):178–92.

81. Lopez J V., Cevario S, O’Brien SJ. Complete nucleotide sequences of the domestic cat (Felis catus) mitochondrial genome and a transposed mtDNA tandem repeat (Numt) in the nuclear genome. Genomics [Internet]. 1996 Apr 15;33(2):229–46. Available from: http://www.ncbi.nlm.nih.gov/pubmed/8660972

82. Lopez J V, Yuhki N, Masuda R, Modi W, Brien SJO. Numt, a Recent Transfer and Tandem Amplification of Mitochondrial DNA to the Nuclear G e n o m e of the Domestic Cat. J Mol Evol. 1994;39:174–90.

83. Antunes A, Ramos MJ. Discovery of a large number of previously unrecognized mitochondrial pseudogenes in fish genomes. Genomics. 2005;86(6):708–17.

84. Jae-Heup K, Eizirik E, O’Brien SJ, Johnson WE. Structure and patterns of sequence variation in the mitochondrial DNA control region of the great cats. Mitochondrion [Internet]. 2001 Oct;1(3):279–92. Available from: http://www.ncbi.nlm.nih.gov/pubmed/16120284

85. Kim J-H, Antunes A, Luoa S-J, Menninger J, Nash WG, O’Brien SJ, et al. Evolutionary analysis of a large mtDNA translocation (numt) into the nuclear genome of the Panthera genus species. Brain, Behav Immun. 2006 Jul;22(5):629–629.

86. Armstrong EE, Taylor RW, Miller DE, Kaelin C, Barsh G, Hadly EA, et al. Long live the king: chromosome-level assembly of the lion (Panthera leo) using linked-read, Hi-C, and long read data. BMC Biol [Internet]. 2020;18(3):705483. Available from: https://www.biorxiv.org/content/10.1101/705483v1.abstract

87. Pritchard JK, Stephens M, Donnelly P. Inference of population structure using multilocus genotype data. Genetics [Internet]. 2000 Jun;155(2):945–59. Available from: http://www.pubmedcentral.nih.gov/articlerender.fcgi?artid=1461096&tool=pmcentrez&rendertype=abstract

88. Evanno G, Regnaut S, Goudet J. Detecting the number of clusters of individuals using the software STRUCTURE: a simulation study. Mol Ecol [Internet]. 2005 Jul [cited 2012 Mar 9];14(8):2611–20. Available from: http://www.ncbi.nlm.nih.gov/pubmed/15969739

89. Earl DA, VonHoldt BM. STRUCTURE HARVESTER: a website and program for visualizing STRUCTURE output and implementing the Evanno method. Conserv Genet Resour [Internet]. 2012 Oct 13 [cited 2014 Jul 9];4(2):359–61. Available from: http://link.springer.com/10.1007/s12686-011-9548-7

90. Jakobsson M, Rosenberg NA. CLUMPP: a cluster matching and permutation program for dealing with label switching and multimodality in analysis of population structure. Bioinformatics [Internet]. 2007 Jul 15 [cited 2014 Jan 10];23(14):1801–6. Available from: http://www.ncbi.nlm.nih.gov/pubmed/17485429

91. Peakall R, Smouse PE. GenAlEx 6.5: genetic analysis in Excel. Population genetic software for teaching and research--an update. Bioinformatics [Internet]. 2012 Oct 1 [cited 2013 Dec 11];28(19):2537–9. Available from: http://www.pubmedcentral.nih.gov/articlerender.fcgi?artid=3463245&tool=pmcentrez&rendertype=abstract

92. Petkova D, Novembre J, Stephens M. Visualizing spatial population structure with estimated effective migration surfaces. Nat Genet. 2016;48(1):94–100.

93. IUCN SSC Cat Specialist Group. Conservation strategy for the lion in West and Central Africa. Gland, Switzerland: IUCN; 2006. 1–60 p.

