## Supplemental Information S1 for "Whole genome sequencing and the application of a SNP panel reveal primary evolutionary lineages and genomic variation in the lion (*Panthera leo*)"

**Supplemental Information S1:**
**Details on laboratory protocols, sequencing, mapping, SNP calling and quality control**

#### Laboratory protocols and sequencing

DNA was extracted using the DNeasy Blood & Tissue kit (Qiagen) following the manufacturer's protocol. The DNA of ten lions and one leopard was sequenced on 3 lanes of an Illumina HiSeq2000 to 99 bp paired end reads with 200-400 bp insert size (Leiden Genome Technology Center, Leiden, The Netherlands), using the Illumina NEB-Next kit for sample preparation. In the first run, two individuals (Benin and Kenya) were tagged and pooled with leopard DNA as the out-group (ratios 1:1:2 for Benin, Kenya and Leopard respectively, to ensure sufficient coverage for the outgroup species). In the two following runs, we tagged four individuals (Cameroon+Somalia+RSA+India and DRC+Zambia1+Zambia2+Namibia), which were equimolarily pooled (Supplemental Table 1). Resulting datasets were demultiplexed based on the unique barcode sequences. After removing adapter sequences with cutadapt [1] and quality trimming with Sickle [2], quality was assessed using FastQC [3].

#### Quality control

FastQC results of the first Illumina run showed a severe drop in quality in the second read (Supplemental Figure 1, below). Therefore, the first run, containing Benin, Kenya and Leopard was repeated. We hard-clipped the reads of the first run after the first 30 bp and added these data to the reads derived from a second run containing the same samples.

After whole genome sequencing, samples Benin and RSA showed bimodal distributions of GC content per read and high average GC content compared to the other samples (55% and 45%, respectively, versus ~40%), indicating contamination with bacterial DNA (Supplemental Figure 2, below). A nucleotide Blast search [4] was done on a random selection of 10,000 reads per sample. Bacterial genomes of the highest hits were downloaded from GenBank (Supplemental Table 7) and reads for samples Benin and RSA were aligned against these using BWA-MEM [5]. Only unaligned reads were retained. In a second filtering step, only reads aligning to the tiger reference genome [6] were included for downstream analyses. Re-analysis of the GC content distribution for these samples showed that these filtering steps eliminated the second peak (Supplemental Figure 2, below). The total number of reads per sample, before and after filtering, is reported in Supplemental Table 1.

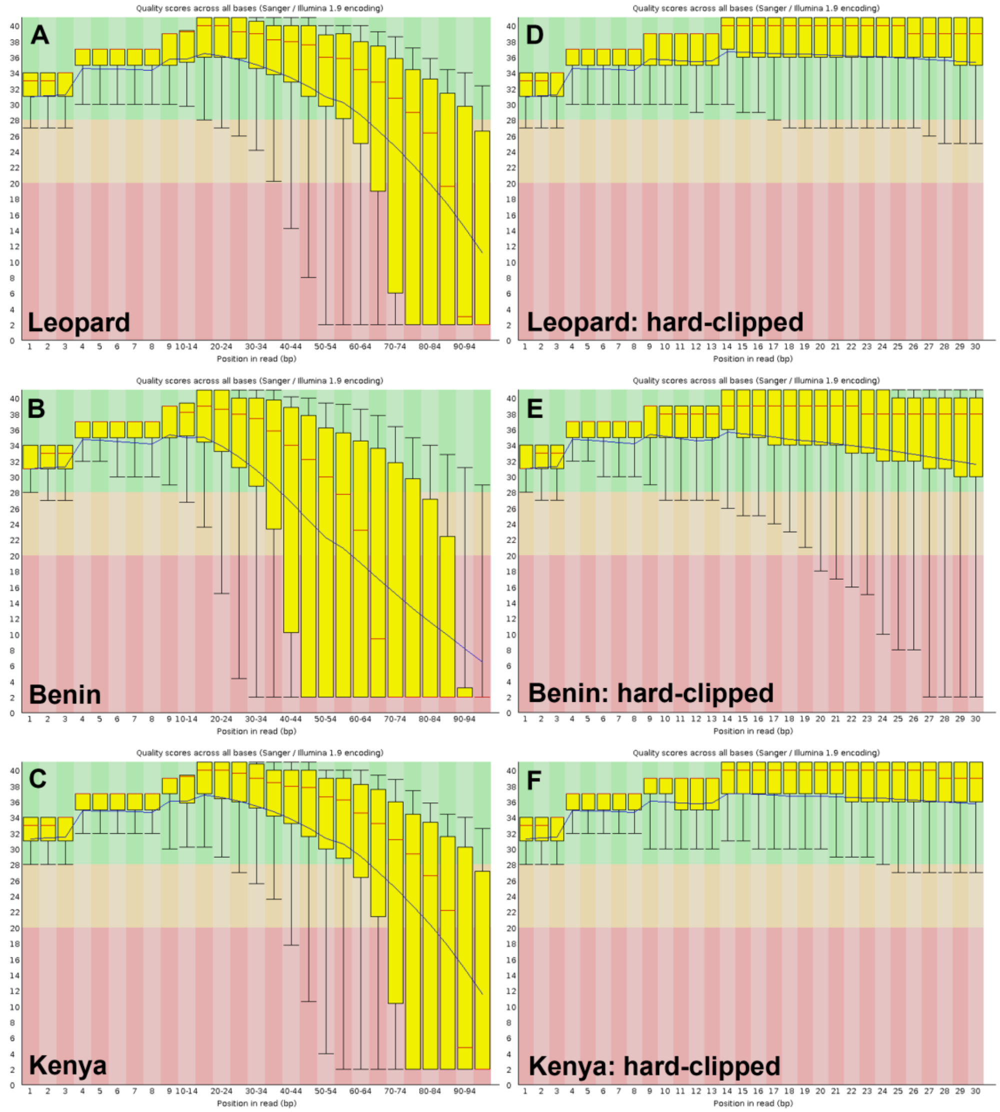


**Supplemental Figure S1. Read quality derived from the first Illumina run, containing one leopard and two lion samples.** Drop in quality scores for (A) Leopard, (B) Benin and (C) Kenya and quality scores after hard clipping of reads after 30 bp for (D) Leopard, (E) Benin and (F) Kenya.

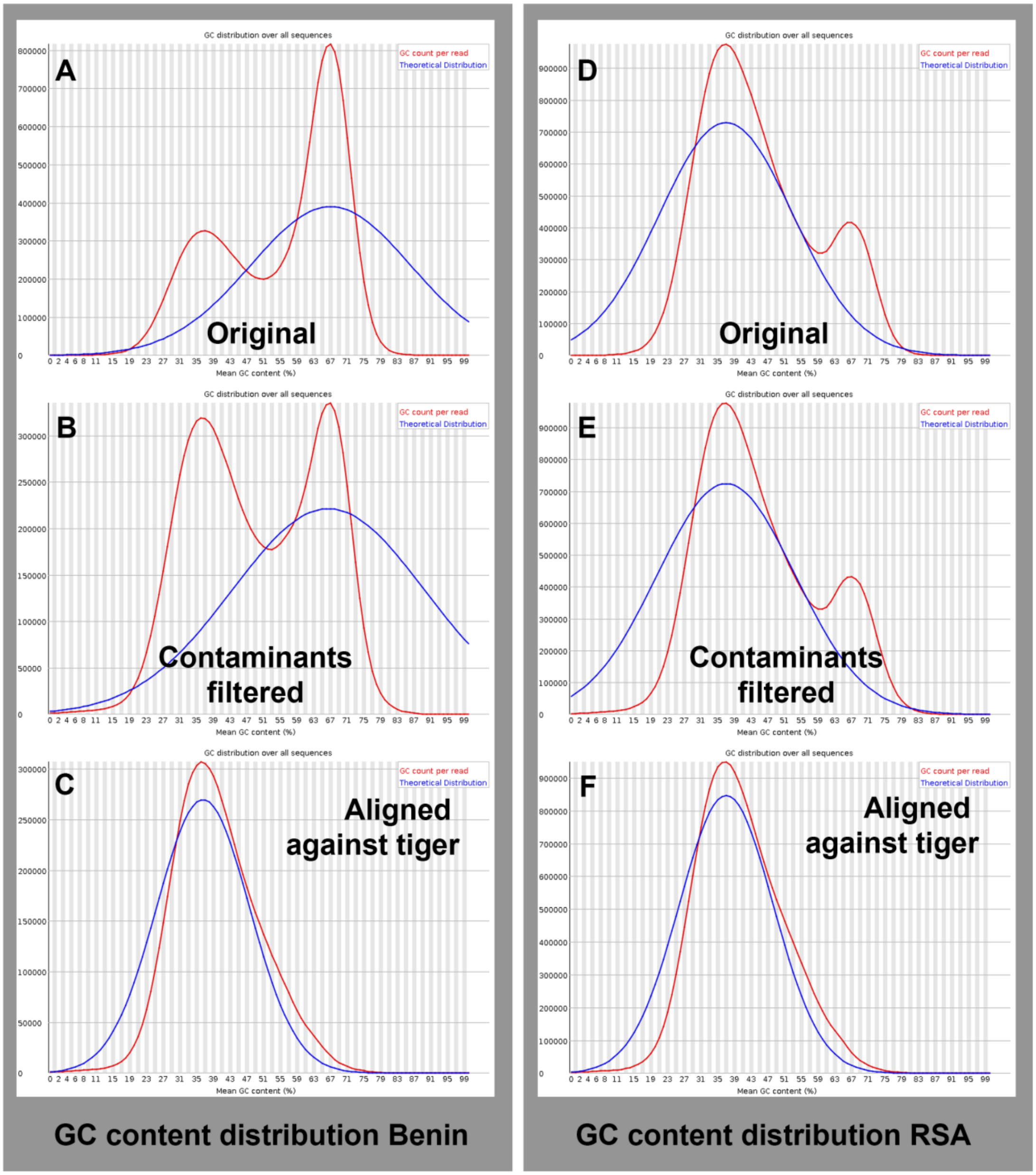


**Supplemental Figure S2. GC content distribution for two lion samples showing signs of bacterial contamination.** GC content of raw reads of (A) Benin and (D) RSA, (B+E) reads filtered against main contaminants and (C+F) reads after filtering and aligned against the tiger reference genome.

**Mapping and SNP calling**

As a reference genome we used the Amur tiger (*Panthera tigris altaica*) autosome assembly [6] (assembly N50 = 8.8 MB) and supplementing this with a lion mitogenome (Genbank Accession number: KP001493). This mitogenome was generated using long range PCR (see [7] for details), therefore minimizing the risk of including nuclear copies of mitochondrial sequences (numts). The known numt sequence was appended to the lion mitogenome reference, enabling filtering of the reads which map to the true mtDNA sequence and reads which map to the numt. Reads of all lions and Leopard were aligned to this reference using BWA-MEM [5]. Reads of all samples were sorted with SAMtools [8] and duplicates were flagged and removed using Picard (http://broadinstitute.github.io/picard, v.1.130). Mapping of the reads produced an average coverage of 3.8 for the full lion genomes (36x for the Leopard genome), ranging from 1.6x (Benin) to 6.8 (Cameroon) (Supplemental Table 1). Only reads mapping to the mitogenome were kept for the mitogenome alignment (i.e., reads mapping to the appended numt sequence were clipped). For the mitogenome we obtained an average coverage of 258.1x for the lion genomes (829.3x for the Leopard genome) (Supplemental Table 1).

Single Nucleotide Polymorphism (SNP) calling was performed using two approaches: 1) using genotype likelihoods as is implemented in ANGSD [9], and 2) using genotype calls from the HaplotypeCaller in GATK [10]. We ran ANGSD (-minQ 20 -GL 2 -doCounts 1 -setMinDepth 27 -setMaxDepth 80 -doMajorMinor 1 -doMaf 1 -SNP_pval 2e-6) and we explored using different numbers of individuals for which data were available at each position (-minInd flag, ranging from 3 to 10). For the SNP genotypes from GATK, we merged the resulting gVCF files with GenotypegVCF, implemented in GATK, as it is suggested as the best practices work flow by the developers. Resulting VCF files were further analyzed using VCFtools [11]. SNPs were called by selecting di-allelic, variable positions within the ten lion samples, filtered for indels and minimum quality (phred score ≥ 20). In addition, we filtered for coverage by selecting the sites with mean depth (over all included individuals) being minimally 3 and maximally 8 (lion samples were pooled aiming for an average coverage of 3-4). Because of the low sequencing depth of the Benin sample as a result of contamination (see 'Quality control'), this individual was excluded from the calculation of mean depth. The resulting VCF file was supplemented with base calls for the samples Benin and the outgroup (Leopard) and filtered for coverage on the individual level, in which we only retained SNPs with a minimum coverage of 3 and replacing positions with lower coverage with ambiguous nucleotides (N). Full mitogenomes were derived for all included samples and were previously reported in [7], clipping all reads which mapped to the numt sequence. Calling Y chromosomal SNPs in the eight samples from male lions was done on scaffolds supposedly located on the Y chromosome, identified by aligning all known Y chromosomal regions in cat (*Felis catus*) to the annotated tiger genome from Cho *et al.* (2013) (Supplemental Table 8). For Y-chromosomal SNPs, we configured SAMtools mpileup to assume a haploid genome; all positions with a heterozygote calling (22 out of 164) were discarded. Coverage plots for Y-chromosomal scaffolds are shown in Supplemental Figure 3 (below).

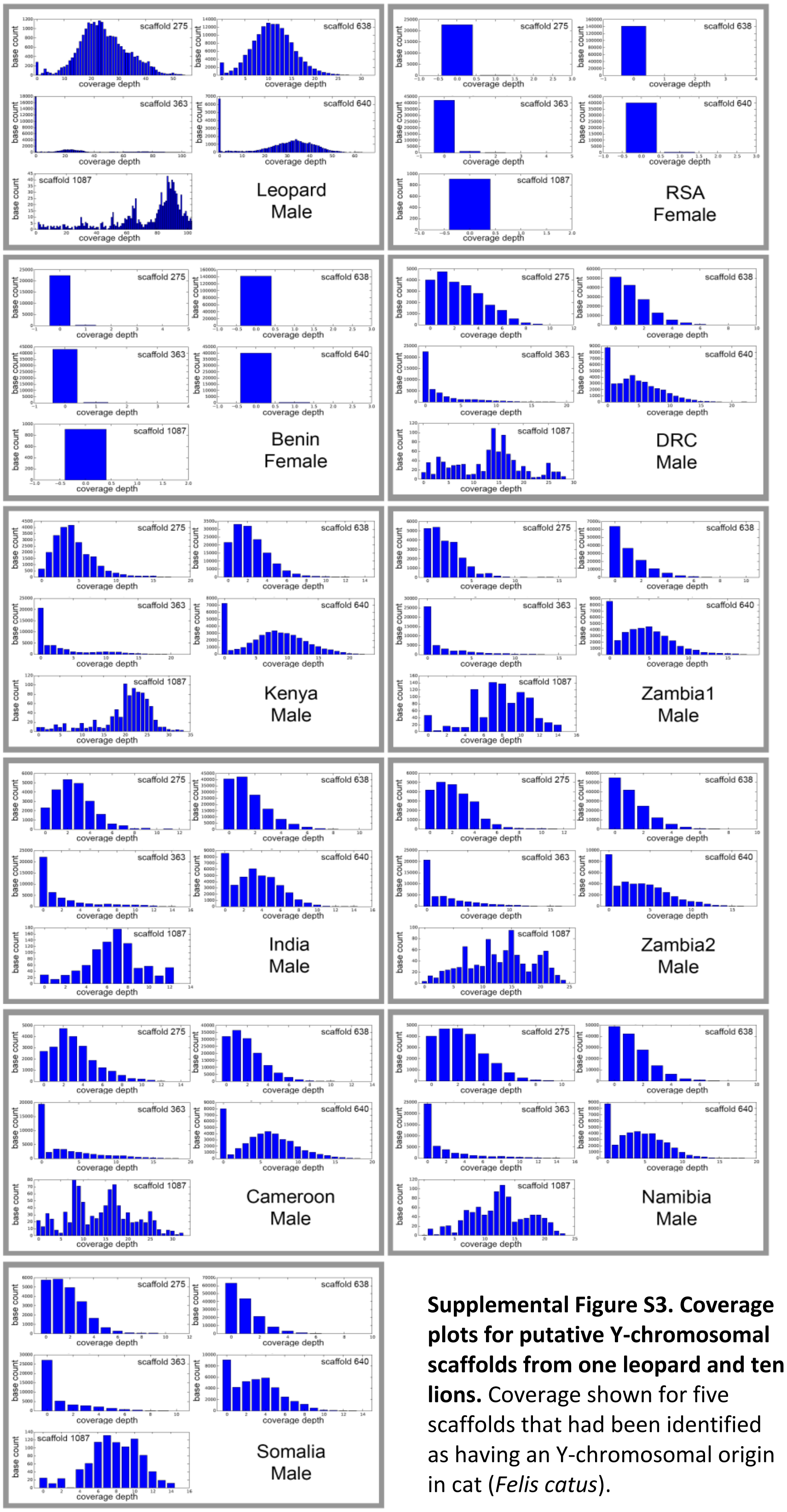


**Supplemental Figure S3. Coverage plots for putative Y-chromosomal scaffolds from one leopard and ten lions.** Coverage shown for five scaffolds that had been identified as having an Y-chromosomal origin in cat (*Felis catus*).

We created different SNP datasets depending on the level of filtering: 1) all SNPs, 2) SNPs called in at least three samples, 3) SNPs called in at least five samples, 4) SNPs called in at least eight samples, and 5) SNPs called in all samples (Supplemental Table 2). Assuming identical chromosomal architecture in the lion and in the tiger, we found a strong relationship between discovered SNPs in this study and estimated chromosome sizes in the tiger [6] (Supplemental Table 2). On the putative Y-chromosomal reference scaffolds, we identified a total of 21 SNPs. Coverage plots for all individuals and all scaffolds confirm the Y-chromosomal origin, as hardly any coverage is found for the two female samples (Supplemental Figure 3, above). Due to low coverage, this Y-chromosomal alignment was not further subjected to phylogenetic analyses. Mitochondrial genomes, consisting of 16,756 bp, excluding repetitive regions RS-2 and RS-3 [12] as was the case in previous phylogeographic analyses [7], were added to the dataset. Mitochondrial and autosomal SNP data (both genotype likelihoods and called genotypes) were subjected to downstream analyses, see main text and Supplemental Information 2.

**SNP panel**

Based on the discovered SNPs, a SNP panel was designed to genotype a larger numbers of samples (see main text). A total of 240 SNPs were selected based on criteria as listed in the main text. After a test run including 10 samples from different popuatlions and with different starting concentrations of DNA, we retained 146 SNPs based on the positions with the highest genotyping success. In addition, primers were designed for 14 mtDNA SNPs, all of which were retained in the final SNP panel. A total of 211 samples were selected for genotyping with the SNP panel. DNA was extracted using the DNeasy Blood & Tissue kit (Qiagen) following the manufacturer's protocol and a Whole Genome Amplification (WGA) kit (LGC genomics) was used to increase the starting amount of DNA. Samples were genotyped at the SNP genotyping facility of the Institute of Biology, Leiden University, using the KASP technique (LGC Genomics) which uses allele specific primers to determine the genotype. Results were analysed in Kraken (LGC Genomics) and manually checked. A pilot study had shown that called genotypes remain consistent with and without the use of a WGA kit (results not shown). In addition, we explored different concentrations of the starting amount of DNA by testing 4 different dilutions of samples, and we included a number of samples which were known to be of lower quality DNA (older, degraded samples). We observed an increase in missing values for samples of lower quality or quantity of DNA (including higher dilutions), however, we never observed changes in the called genotype (allelic drop-out). This illustrates that the SNP panel is producing reliable results, although the sensitivity of downstream analyses with high numbers of missing values needs to be taken into account. Samples showing a high level of missing data (>50%) were included in a subsequent run and results were combined. For the final dataset, the number of missing values was calculated per individual.

Further selection of SNP panel SNPs was performed after genotyping 211 samples for all 146 autosomal SNPs and 14 mitochondrial SNPs. After testing for Hardy-Weinberg equilibrium in multiple samples from seven wild reference populations (Benin, Cameroon, DRC, Kenya, Zambia, Zimbabwe, India), we excluded 7 loci which showed a significant (>5%) deviation from the Hardy-Weinberg equilibrium in all of the 7 tested populations. We further excluded 5 loci with >30% missing data in the entire dataset. We retained 125 autosomal SNPs in the SNP panel, i.e. we excluded 21 SNPs. Positions which showed a conflict after genotyping a sample twice (if the sample had >50% missing data after the first run), as well as heterozygotes in mtSNPs (which are expected to be haploid) were replaced by ambiguous nucleotides (hereafter: missing values).

Since the SNPs originally were selected following the genomic architecture of the tiger genome (to ensure good coverage across the chromosomes), we used BLASTn to find associated coordinates in the recently publishes lion genome [13], which was not yet available at the time of the original mapping.

Coordinates in both tiger and lion genome are reported in Supplemental Table 3 and an overview of the SNP panel SNPs in the lion genome is shown in Supplemental Figure 4 (below).

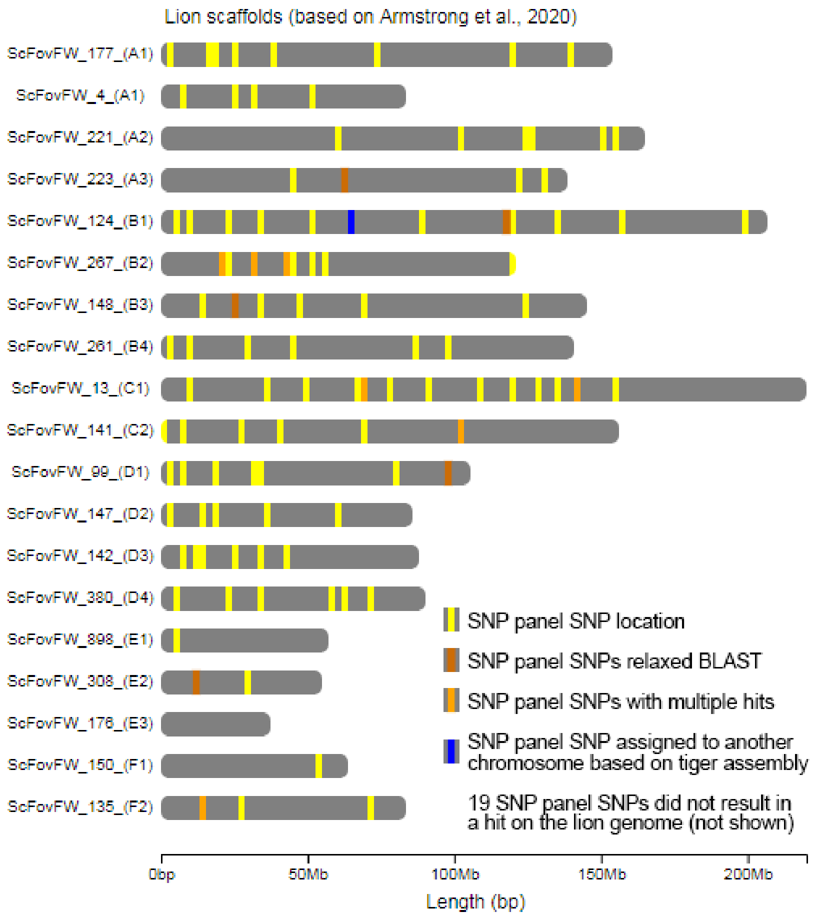


**Supplemental Figure S4. Visualization of the putative localities of the SNP panel SNPs on the lion genome.** Yellow lines indicate localities of SNP panel SNPs; dark orange indicates a hit with relaxed BLAST settings (-word_size 20, both primers found); orange indicates multiple hits on the lion genome (coordinates for lowest E-value shown); blue indicates a SNP which is mapped to a different chromosome depending in the tiger reference. Note that two scaffolds refer to chromosome A1.

### References


1. Martin M (2011) Cutadapt removes adapter sequences from high-throughput sequencing reads. EBMnet.journal 17: 10–12.
2. Joshi NA, Fass JN (2011) Sickle: A sliding-window, adaptive, quality-based trimming tool for FastQ files. Available: https://github.com/najoshi/sickle.
3. Andrews S (2010) FastQC - A Quality Control tool for High Throughput Sequence Data. Available: http://www.bioinformatics.babraham.ac.uk/projects/fastqc/.
4. Altschul SF, Gish W, Miller W, Myers EW, Lipman DJ (1990) Basic Local Alignment Search Tool Department of Computer Science. J Mol Biol 215: 403–410.
5. Li H, Durbin R (2009) Fast and accurate short read alignment with Burrows-Wheeler transform. Bioinformatics 25: 1754–1760. Available: http://www.pubmedcentral.nih.gov/articlerender.fcgi?artid=2705234&tool=pmcentrez&rendertype=abstract. Accessed 9 July 2014.
6. Cho YS, Hu L, Hou H, Lee H, Xu J, et al. (2013) The tiger genome and comparative analysis with lion and snow leopard genomes. Nat Commun 4. doi:10.1038/ncomms3433.
7. Bertola LD, Jongbloed H, Van Der Gaag KJ, De Knijff P, Yamaguchi N, et al. (2016) Phylogeographic Patterns in Africa and High Resolution Delineation of Genetic Clades in the Lion (Panthera leo). Sci Rep 6: 1–11. Available: http://dx.doi.org/10.1038/srep30807.
8. Li H, Handsaker B, Wysoker A, Fennell T, Ruan J, et al. (2009) The Sequence Alignment/Map format and SAMtools. Bioinformatics 25: 2078–2079. Available: http://www.pubmedcentral.nih.gov/articlerender.fcgi?artid=2723002&tool=pmcentrez&rendertype=abstract. Accessed 9 July 2014.
9. Korneliussen TS, Albrechtsen A, Nielsen R (2014) ANGSD: Analysis of Next Generation Sequencing Data. BMC Bioinformatics 15: 1–13. doi:10.1186/s12859-014-0356-4.
10. McKenna A, Hanna M, Banks E, Sivachenko A, Cibulskis K, et al. (2010) The Genome Analysis Toolkit: A MapReduce framework for analyzing next-generation DNA sequencing data. Genome Res 20: 1297–1303. doi:10.1101/gr.107524.110.20.
11. Danecek P, Auton A, Abecasis G, Albers CA, Banks E, et al. (2011) The variant call format and VCFtools. Bioinformatics 27: 2156–2158. doi:10.1093/bioinformatics/btr330.
12. Jae-Heup K, Eizirik E, O'Brien SJ, Johnson WE (2001) Structure and patterns of sequence variation in the mitochondrial DNA control region of the great cats. Mitochondrion 1: 279–292. Available: http://www.ncbi.nlm.nih.gov/pubmed/16120284.
13. Armstrong EE, Taylor RW, Miller DE, Kaelin C, Barsh G, et al. (2020) Long live the king: chromosome-level assembly of the lion (Panthera leo) using linked-read, Hi-C, and long read data. BMC Biol 18: 705483. Available: https://www.biorxiv.org/content/10.1101/705483v1.abstract.
