## Supplemental Information S2 for "Whole genome sequencing and the application of a SNP panel reveal primary evolutionary lineages and genomic variation in the lion (*Panthera leo*)"

**Supplemental Information 2:**
**Validation of phylogenetic inferences and populations structure from whole genome sequencing and SNP panel data**

**Whole genome SNPs**
As individual coverage is relatively low, we used both genotype likeihoods (derived from ANGSD) and genotype calls (using GATK) to infer population structure and phylogenetic relationships. We use a comparison between both methods to further strengthen and validate the results presented in this manuscript.
A dendrogram (Supplemental Figure 5, below) based on genotype likelihoods from all 10 lions (using hclust) shows similar relationships as inferred from Bayesian, Maximum Likelihood and Quartet analyses of genotypes (Figure 1, main text). A basal split is found between West & Central African populations and populations from East and Southern Africa. In the latter, we observe a split between East Africa and Southern Africa, as is also observed from the SNP panel data (Figure 2C, main text). The position of the India sample is likely to be strongly affected by the extremely low genetic diversity in this population, hence affecting its positioning in the dendrogram.

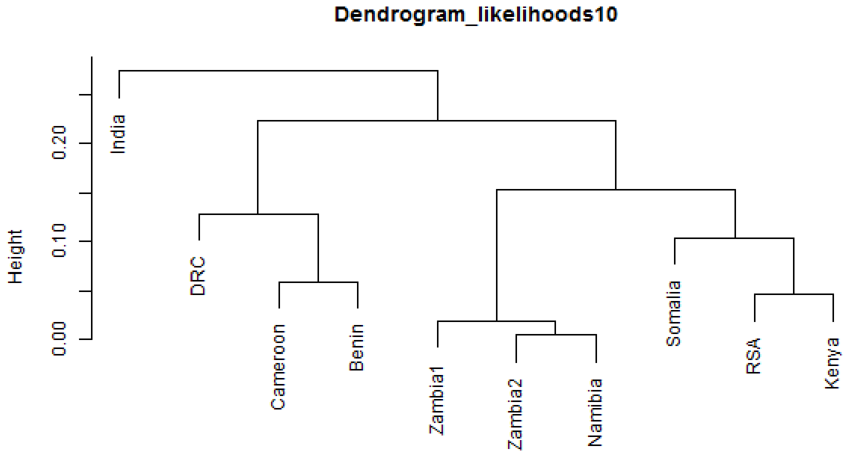


**Supplemental Figure S5. Dendrogram based on the SNP genotype likelihoods, using hclust.**

PCA analysis on SNPs called in all 10 individuals (i.e., no missing data allowed) shows a north-south gradient across PC1, with India and West & Central Africa on one side, and the East & Southern African populations on the other (Supplemental Figure 6, below).

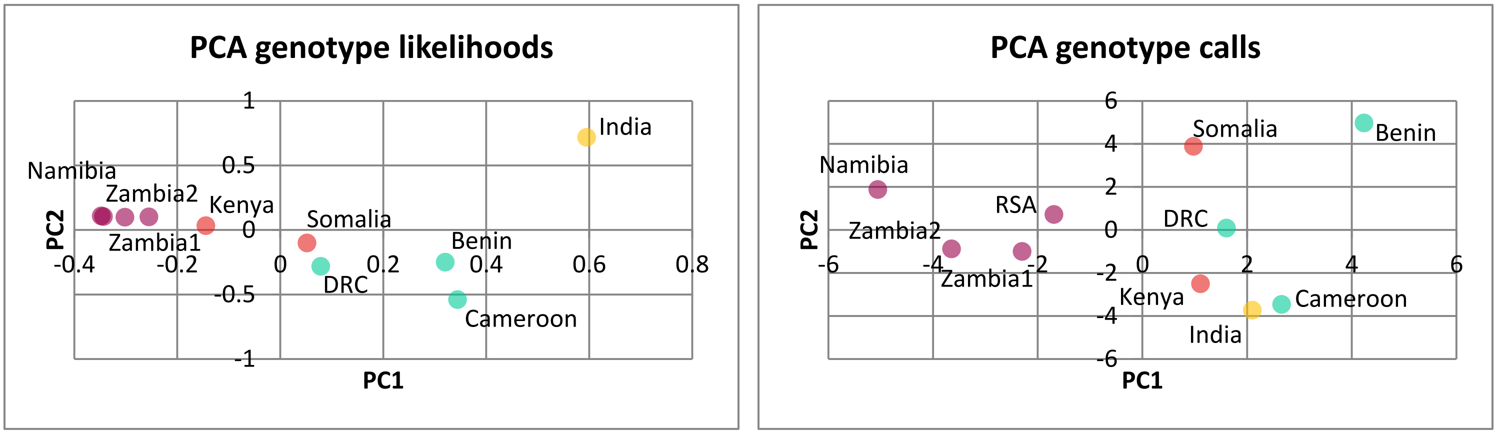


**Supplemental Figure S6. PCA based on genotype likelihoods from ANGSD (left, also included as Figure 3A in main text) and genotypes from GATK (right).** Green refers to West and Central Africa (Benin, Cameroon, DRC), yellow refers to India, red refers to East Africa (Kenya, Somalia), purple refers to Southern Africa (Zambia1, Zambia2, RSA, Namibia).

To assess population structure and possible admixture, we inferred ancestry coefficients from NGSadmix, based on genotype likelihoods, and from sNMF (LEA package), based on genotype calls (Supplemental Figure 7, below). The results from the latter were used to project inferred population structure on a map.

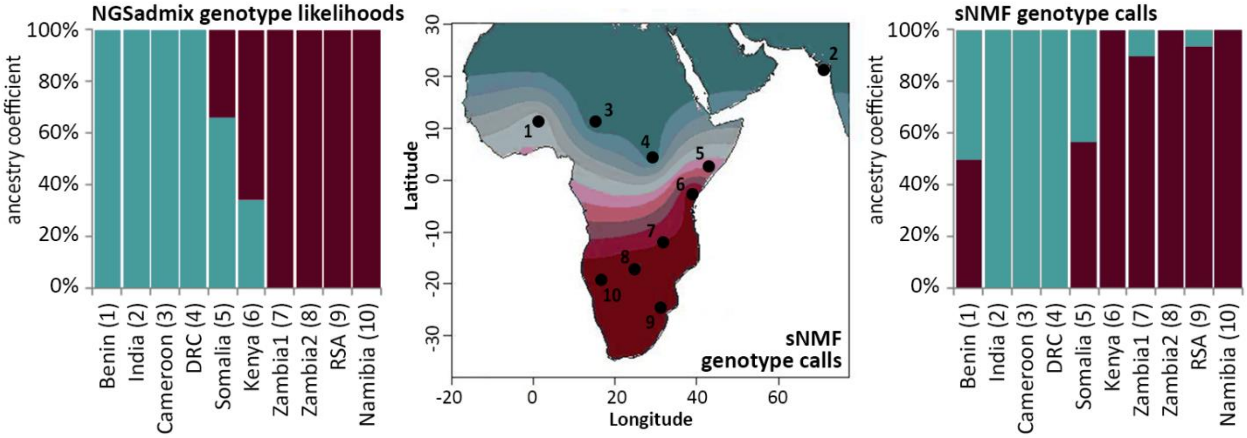


**Supplemental Figure S7. Comparison ancestry coefficients genotype likelihoods using NGSadmix (left, also included as Figure 2A in main text) and derived from SNP genotypes using sNMF (right).** A spatial representation of ancestry coefficients from sNMF is shown as a map (middle).

Although sNMF results seem to suggest a mixed origin for the sample from Benin (sample 1), we hypothesize that this pattern is an artefact attributable to the very high number of missing data from this sample. This particular pattern is not visible in the NGSadmix results, in which Benin clusters strongly to other populations from the *P. leo leo* subspecies. In addition, the PCA plots in which no missing data were allowed, also show that Benin does not have an intermediate position between *P. leo leo* and East African populations. Finally, this is further corroborated by SNP panel results, in which no pattern of admixture is detected in the five included samples from Benin.

Both NGSadmix and sNMF suggest admixture in East Africa, where both subspecies overlap and interbreed. Both methods suggest admixed origin for Somalia and NGSadmix suggests a lower degree of admixture in Kenya, which is not unlikely due to the geographic location of this population. This is corroborated by using D-statistics and the abbababa methods as implemented in ANGSD. More details regarding putative admixture are reported in the main text.

**SNP panel SNPs**

As we are dealing with samples of varying quality and some of the applied methods are sensitive to missing data, we explore the generated datasets on two levels: 1) the complete dataset, and 2) the dataset excluding samples with >25% missing data (results shown in the main text). The number of missing values for the autosomal SNPs and the mitochondrial SNPs, are available in Supplemental Table 4 and summarized in Suppplemental Figure 8 (below).

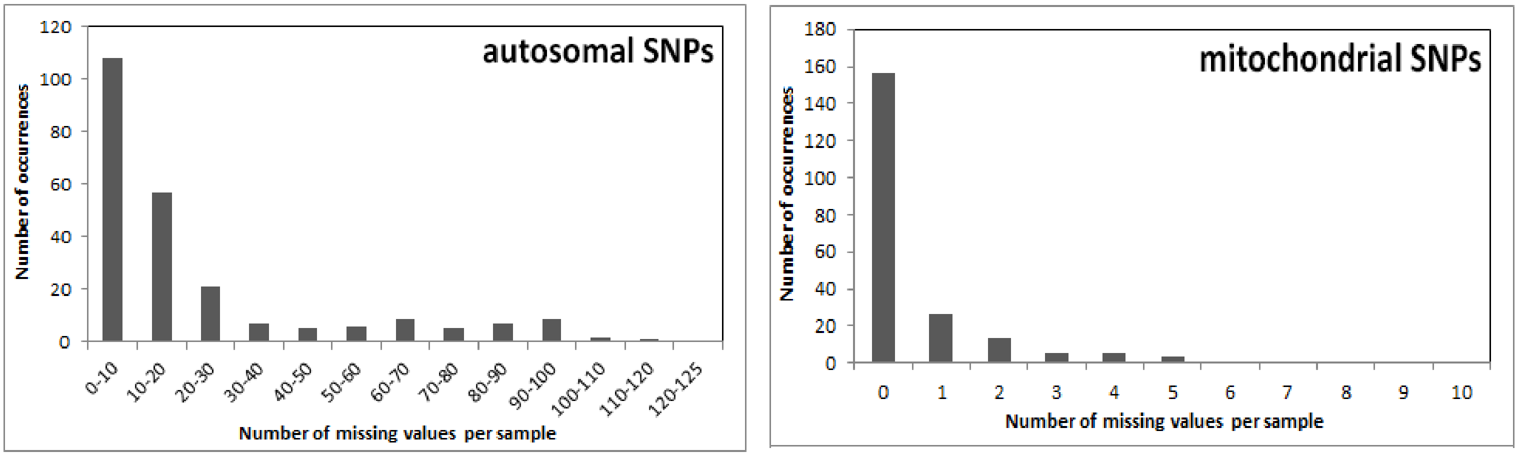


**Supplemental Figure S8. Distribution of missing data in 211 lions which were genotyped using 125 autosomal and 14 mitochondrial SNPs.**

Comparing the results of STRUCTURE when including all samples (i.e., including samples with >25% missing data) (Supplemental Figure 9, below) to the results presented in the main text (Figure 2), several of the Kenyan individuals show assignment values to West and Central Africa. These assignments are likely an artefact of missing data in these individuals,  since this pattern in not apparent in Figure 2 and comparison with mtDNA SNPs does not suggest an association of these individuals to the West & Central African lineage. It must be noted that when K=2, due to the very low diversity in the Indian population, assignment values are driven to ~1 in the Indian lions. Subsequently, samples from West and Central Africa are assigned more strongly to the cluster representing the southern subspecies, with only weak assignment to the India cluster. We emphasize that this does not indicate region-wide hybridization between the two subspecies, nor does it imply that the populations with the highest assignment values constitute ancestral populations. Rather, this result is driven by the extremely low genetic diversity in the Indian population, and is comparable to the Africa/India split which is found based on STRUCTURE analysis or PCA of microsatellite data (Bertola *et al.* 2015), that similarly leads to the strong assignment of the Indian population as a separate cluster. PCA based on SNP data show a similar pattern, with the variation between African populations only becoming apparent after exclusion of the Indian population (Supplemental Figure 10, below). Distinction between African population becomes more pronounced after excluding individuals with >25% missing data (Supplemental Figure 10, below).

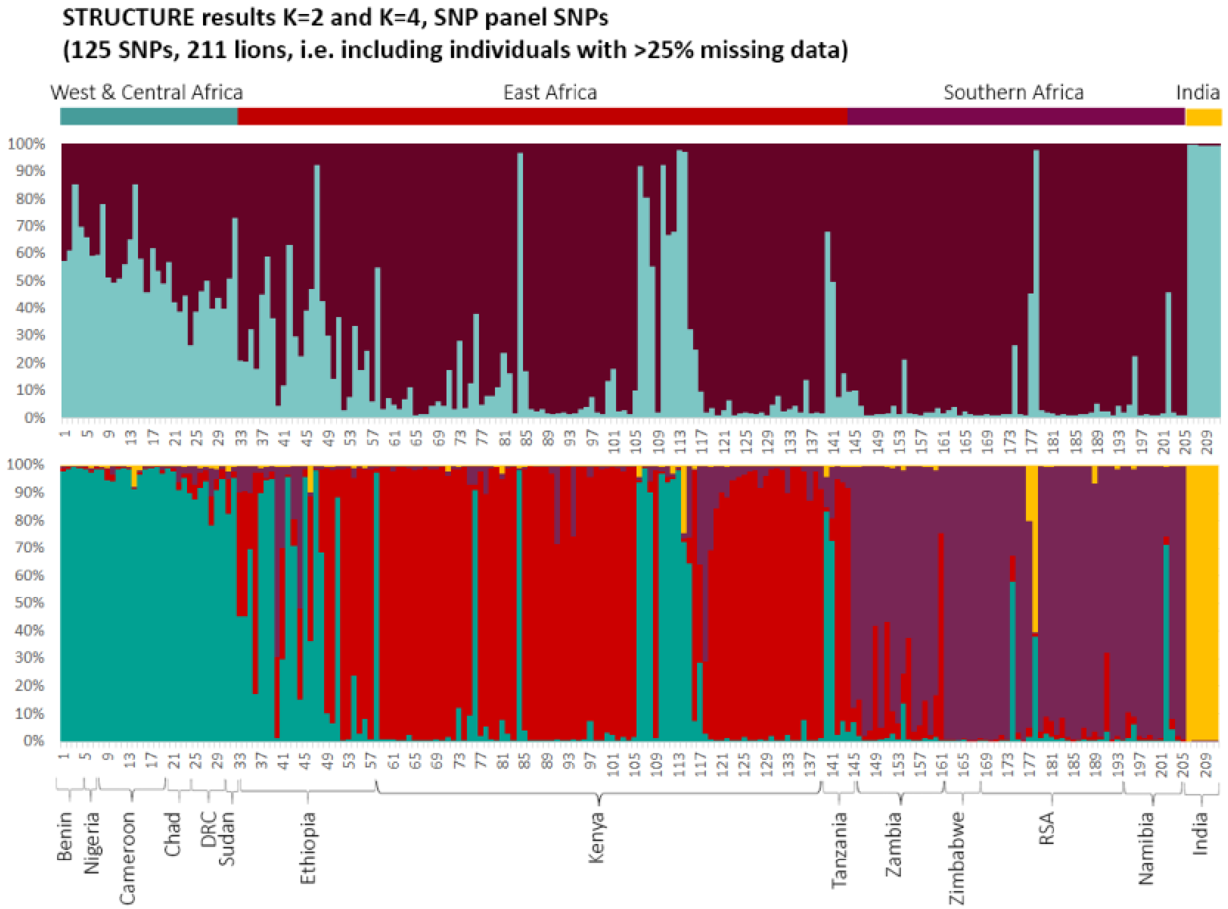


**Supplemental Figure S9. Bar plots indicating assignment values for K=2 and K=4 from a STRUCTURE run, including all samples (211 samples and 125 autosomal SNPs).**

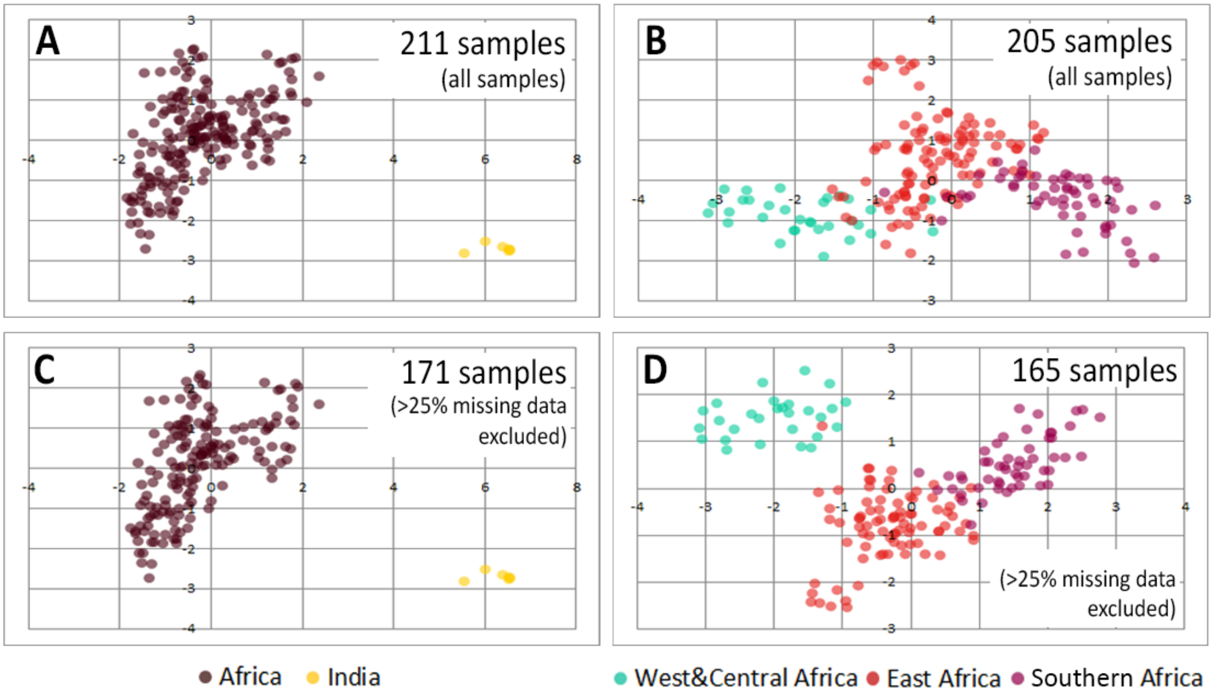


**Supplemental Figure S10. PCA plots based on 125 SNPs in 211 lion samples (A), and after exclusion of the Asiatic population (B).** Panels showing results for 171 samples (i.e. samples with >25% missing data excluded) are also included in the main text as Figure 3B and 3C.

Comparing the assignments based on STRUCTURE to the assignments based on the mitochondrial SNPs show a largely congruent pattern (Figure 2C, Supplemental Table 5). Six individuals show an unexpected haplotype (i.e., a haplotype not previously documented in the country): Ethiopia13 – West African haplotype, Ethiopia26 – East/Southern African haplotype, Kenya49 – Central African haplotype, Tanzania1 – North East African haplotype, RSA10 – North East African haplotype, Namibia2 – North East African haplotype. However, it must be noted that all of these individuals have >35% missing autosomal data (with Kenya49, Tanzania1, RSA10 and Namibia2 even missing >50% autosomal data). In addition, Ethiopia26, RSA10 and Namibia2 also have >20% missing data in the mitochondrial SNPs, and therefore results should be interpreted with caution.
